# A disease mutation reveals a non-swapped pore and rewired gating cycle of the mechanically activated PIEZO1 channel

**DOI:** 10.1101/2025.11.17.688653

**Authors:** Chenxi Wang, Ziying Zhang, Xuzhong Yang, Sijia Liu, Jingyi Yuan, Chao Lin, Xiaochun Zhang, Boxue Tian, Xueming Li, Bailong Xiao

## Abstract

PIEZO1 is a versatile mechanotranduction channel critical for diverse cellular and physiological processes^1–3^. Gain-of-function mutations, such as R2482H, which slow channel inactivation, are linked to dehydrated hereditary stomatocytosis (DHS) and iron overload^4–7^. PIEZO1 forms a gigantic bowl-shaped trimer comprising flattenable peripheral blades and a central pore module with a switchable cap^8–12^. However, the structural dynamics governing its gating process and pathogenesis remain incompletely understood. Here, we determined distinct conformational states of the DHS-associated PIEZO1-R2482H mutant in both detergents and proteoliposomes. Remarkably, the R2482H mutation in the central pore induces a conformational change that propagates ∼25 nm to the most distal blade, enabling resolution of all 38 transmembrane helices. PIEZO1-R2482H is inherently flatter than PIEZO1, lowering the gating force and energy barrier and thus explaining its heightened mechanosensitivity. While the flattened PIEZO1 has its cap in the up-state adopting an inactivated state, the flattened PIEZO1-R2482H has its cap in the down-state and twisted by ∼75°, drastically switching the pore module from a swapped to a non-swapped organization. Such twisted conformation of PIEZO1-R2482H might hinder the transition to the inactivated state, instead proceeding through a deactivation-transition state characterized by curved blades and a partially twisted cap, thereby accounting for the slowed inactivation and deactivation kinetics. Taken together, we reveal the full transmembrane topology and a non-swapped pore conformation induced by R2482H, and propose the structural dynamic model that delineates the distinct gating cycle of activation, inactivation, and deactivation for PIEZO1 and PIEZO1-R2482H.

## Main

The PIEZO family of mechanically activated ion channels, including PIEZO1 and PIEZO2 in mammals, mediate a broad spectrum of in vivo mechanotransduction processes, ranging from PIEZO1-dependent homeostatic regulations of various tissues and cell types^6,7,13–21^ to PIEZO2-dependent somatosensation of touch and pain, proprioception, and interoception in primary sensory neurons^22–29^. In response to mechanical stimuli, PIEZO1/2 rapidly open and subsequently inactivate on the millisecond timescale^1^. Such precise gating dynamics quantitatively control cation permeation and consequently ensure proper regulation of the downstream cellular responses such as membrane depolarization and Ca^2+^ signaling. For example, PIEZO1-mediated Ca^2+^ signaling regulates the volume of red blood cells^13^ and the phagocytic activity of macrophages^19^. Human genetic mutations in *PIEZO1* have been associated with dehydrated hereditary stomatocytosis (DHS) and iron overload^4–7^. Human PIEZO1-R2456H and the corresponding murine orthologous PIEZO1-R2482H represent one of the most well-characterized DHS-associated mutations^4^. Compared to wild-type PIEZO1, both hPIEZO1-R2456H and mPIEZO1-R2482H show apparently slowed inactivation kinetics of their mechanically activated currents^4,5,30^. These findings highlight the importance of PIEZO1’s gating kinetics in determining its biological functions and pathophysiological roles.

Detergent solubilized PIEZO1 forms bowl-shaped trimers comprising a central ion-conducting pore with an extracellular cap and three peripheral curved and non-planar blades, each of which contains 26 well-resolved transmembrane helices (TMs) (TM13-TM38) and a featured intracellular beam^31,32^. The N-terminal region containing the predicted TM1-12 remains unresolved, indicating high intrinsic flexibility^8–12,33^. Nanoscopic fluorescence imaging has verified the large degree of flexibility of the distal blades in cells^34^. The pore domain containing the TM37, termed outer helix (OH), the C-terminal extracellular domain (CED), and the TM38, termed inner helix (IH), is structurally swapped into the neighboring blade domain^10,31^. The intrinsically curved structure might represent a closed state, in which the hydrophobic TM pore with a radius of 2-4 Å is suggested to be locally unwetted and therefore energetically unfavored for water and ion permeation^35^. In contrast to the partially resolved structure of PIEZO1, the PIEZO2 structure reveals all 38 TMs, with the N-terminal 36 TMs organized into nine repetitive transmembrane helical units (THU1-9), and the C-terminal TM37 (OH) and TM38 (IH) forming the swapped central pore module^9^. The relative middle-plane height between the most distal blade measured at TM3 of THU1 and the central IH is ∼9.1 nm, and accordingly the three highly curved blades enclose a nano-bowl structure with a calculated curvature radius of ∼12 nm. Furthermore, the IH-enclosed transmembrane pore of PIEZO2 is more constricted than that of PIEZO1, containing both the upper and lower TM gates^9^.

PIEZO1 serves as the prototypic member for studying the structural dynamics and gating mechanisms of PIEZO channels. PIEZO1 reconstituted in lipid bilayers undergoes reversible flattening at biologically relevant forces, as probed by high-speed atomic force microscopy (HS-AFM)^32^. By reconstituting PIEZO1 into liposome vesicles made from the soy total lipid extract, we determined both curved and flattened structures of PIEZO1 in lipid membranes, demonstrating that PIEZO1 can directly sense forces derived solely from the lipid membrane, triggering a conformational switch from the curved to the flattened state^36^. The flattened structure of PIEZO1 likely represents an inactivated state, with the cap in the up-state and the cap-gate closed^36^. In an attempt to capture the open state structure, we electrophysiologically identified the PIEZO1-S2472E mutant with constitutive opening and slowed inactivation kinetics^30^. We then determined the PIEZO1-S2472E-intermediate-open structure with a curvature radius of ∼32 nm, in which the blades are partially flattened, the cap is in the down-state, and both the cap- and TM gates are open, while the intracellular lateral plug gate remains plugged^30^. Further flattening of the blades and levering of the beam has been proposed to displace the lateral-plug-gate, reaching a fully open state. A flattened structure of PIEZO1 derived from cell membrane vesicles, as reported in a recent preprint^37^, is in line with the expected full open state, in which the blades are fully flattened and the cap is in the down-state. Combining analysis of the series of PIEZO1 and PIEZO1-S2472E structures determined in detergent micelles or liposomes, we assigned the close, intermediate-open, open and inactivated states of PIEZO1 based on their characteristic structural features, leading to the proposal of a curvature-based gating model of PIEZO1 ^30^.

Continuing our efforts to elucidate the gating and disease-causing mechanisms of PIEZO channels, here we focused on the DHS-associated PIEZO1-R2482H mutant and obtained the first topologically complete 38-transmembrane-helix (38-TM) structure of PIEZO1 and a novel conformational state, in which the central pore module is switched from a swapped into a non-swapped configuration. By integrating these findings with previously reported structural states of PIEZO1, we propose the distinct structural dynamics underlying the gating process of activation, inactivation, and deactivation for PIEZO1 and R2482H.

### R2482H structure with a fully resolved 38-TM-topology

Following our established procedure for determining the cryo-EM structures of PIEZO1, PIEZO2, PIEZO1.1, and the PIEZO1-S2472E mutant^9,10,30,33^, and as detailed in the Method and illustrated in the single-particle cryo-EM workflow (Extended Data Fig. 1 and 2), we prepared the PIEZO1-R2482H protein solubilized in the detergent glycol-diosgenin (GDN) and determined its structure by cryo-EM. From a total of three datasets of collected micrographs, we reconstructed the three-dimensional (3D) structure of PIEZO1-R2482H at an overall resolution of 4.18 Å, which closely resembles the three-bladed, propeller-shaped PIEZO1 structure with 26 out of 38 TMs resolved per protomer (Extended Data Fig. 1 and 2 and Extended Data Table 1). Noting that the map reconstructed from the first dataset showed clear density at the distal part of the blade (Extended Data Fig. 2), we performed focused refinements on the distal blades and successfully resolved the remaining 12 TMs. We built the structural model of PIEZO1-R2482H from the map and designated it as R2482H-detergent-curved (Fig. 1a-c and Extended Data Fig. 1), to distinguish it from the curved and flattened structures of R2482H derived from proteoliposomes (Fig. 2a,b), which are termed R2482H-liposome-curved (Fig. 2a) and R2482H-liposome-flattened (Fig. 2b), respectively. Similar to the previously determined structure of PIEZO2 (PDB: 6KG7), PIEZO1-R2482H exhibits a 38-TM-topology comprising the first 36 TMs, which are organized into THU1-9 to form the highly curved peripheral blade domain, and the last 2 TMs as the OH and IH, which, together with the extracellular cap domain and the intracellular CTD, trimerize to form the central pore module (Fig. 1a,b). Given that the most distal TM1-TM12 of wild-type PIEZO1 is too flexible to be resolved, the pore-localized R2482H mutation induces a central-to-peripheral allosteric effect all the way to the most distal blade with a calculated distance of ∼25 nm, resulting in stabilization of the flexible TM1-TM12 and enabling determination of the full 38-TM-topology (Fig. 1a-c, and Extended Data Fig. 1h, i).

**Fig. 1.**
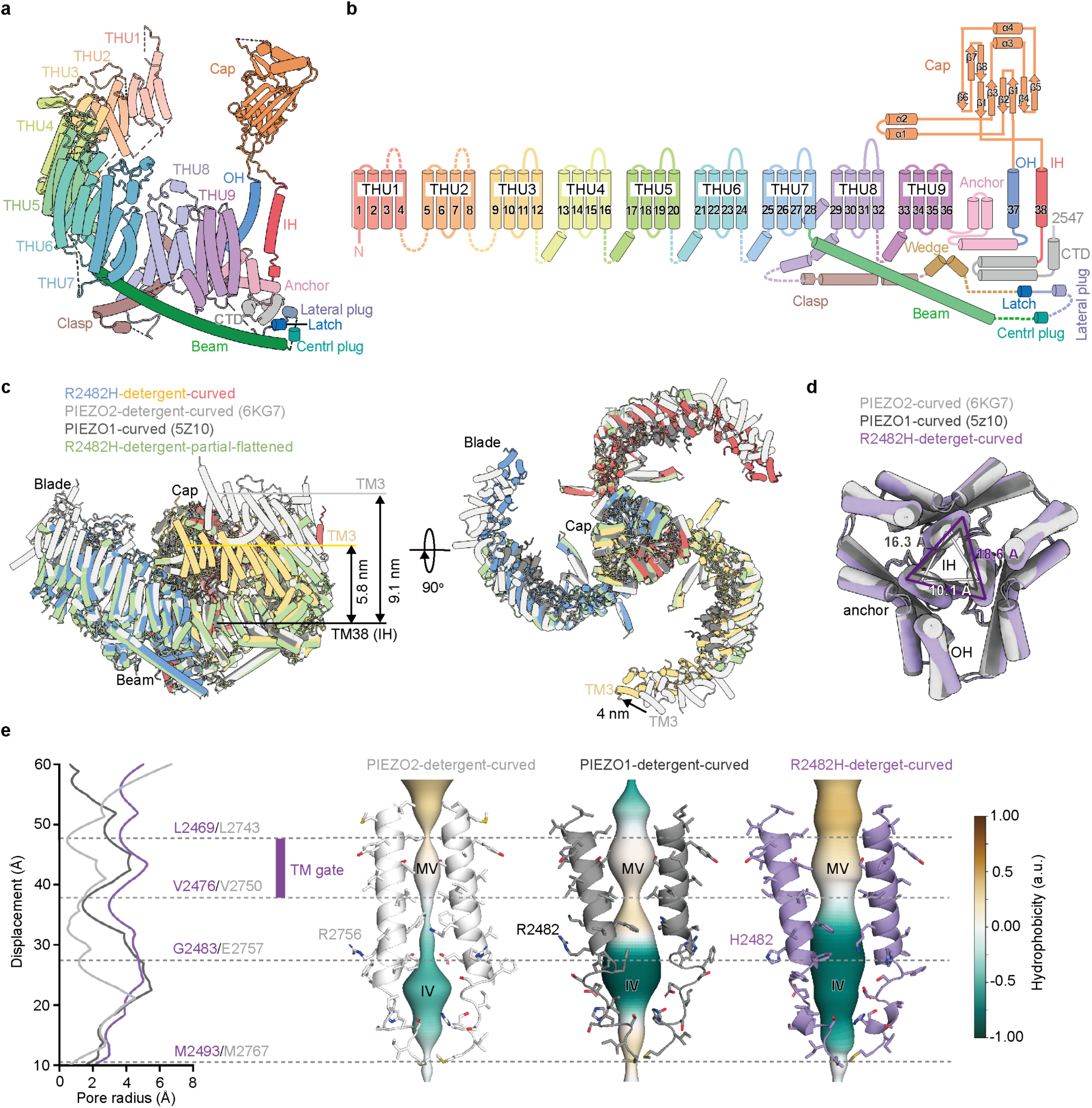
Determination and analyses of PIEZO1-R2482H structures. **a**, The cartoon model of PIEZO1-R2482H protomer showing the indicated structural domains. THU represents the abbreviation of Transmembrane Helical Unit. **b**, The 38-TM-topological model of PIEZO1-R2482H protomer showing the indicated structural domains. The wedge domain is disordered in the curved structure of PIEZO1-R2482H, but becomes ordered in the flattened state shown in Fig. 2. **c**, Side or top views of the superimposed trimeric structures of PIEZO1-detergent-curved (PDB: 5Z10, black), PIEZO2-detergent-curved (PDB: 6KG7, gray), R2482H-detergent-curved (colored), R2482H-detergent-partial-flattened (green). The middle-plane height in between the outmost TM3 of THU1 and the central TM38 (IH) of the full structure of R2482H-detergent-curved and PIEZO2-detergent-curved is 5.8 nm and 9.1 nm, respectively. The lateral displacement of the TM3 of PIEZO2-curved relative to that of R2482H-curved is indicated by the black arrow. **d**, Top view of the superimposed central pore region of PIEZO1-detergent-curved (PDB: 5Z10), R2482H-detergent-curved and PIEZO2-detergent-curved (PDB: 6KG7), showing the distance between two neighboring IHs. **e**, Pore radius and the CHAP-based hydrophobicity profile along the IH-enclosed central axis of PIEZO1-deternget-curved (PDB: 5Z10, black), R2482H-detergent-curved (purple) and PIEZO2-detergent-curved (PDB: 6KG7, gray). The hydrophobic TM gate residues L2469 and V2476 in PIEZO1 and the corresponding residues L2743 and V2750 in PIEZO2 are shown. MV and IV represent the membrane vestibule and intracellular vestibule, respectively.

**Fig. 2.**
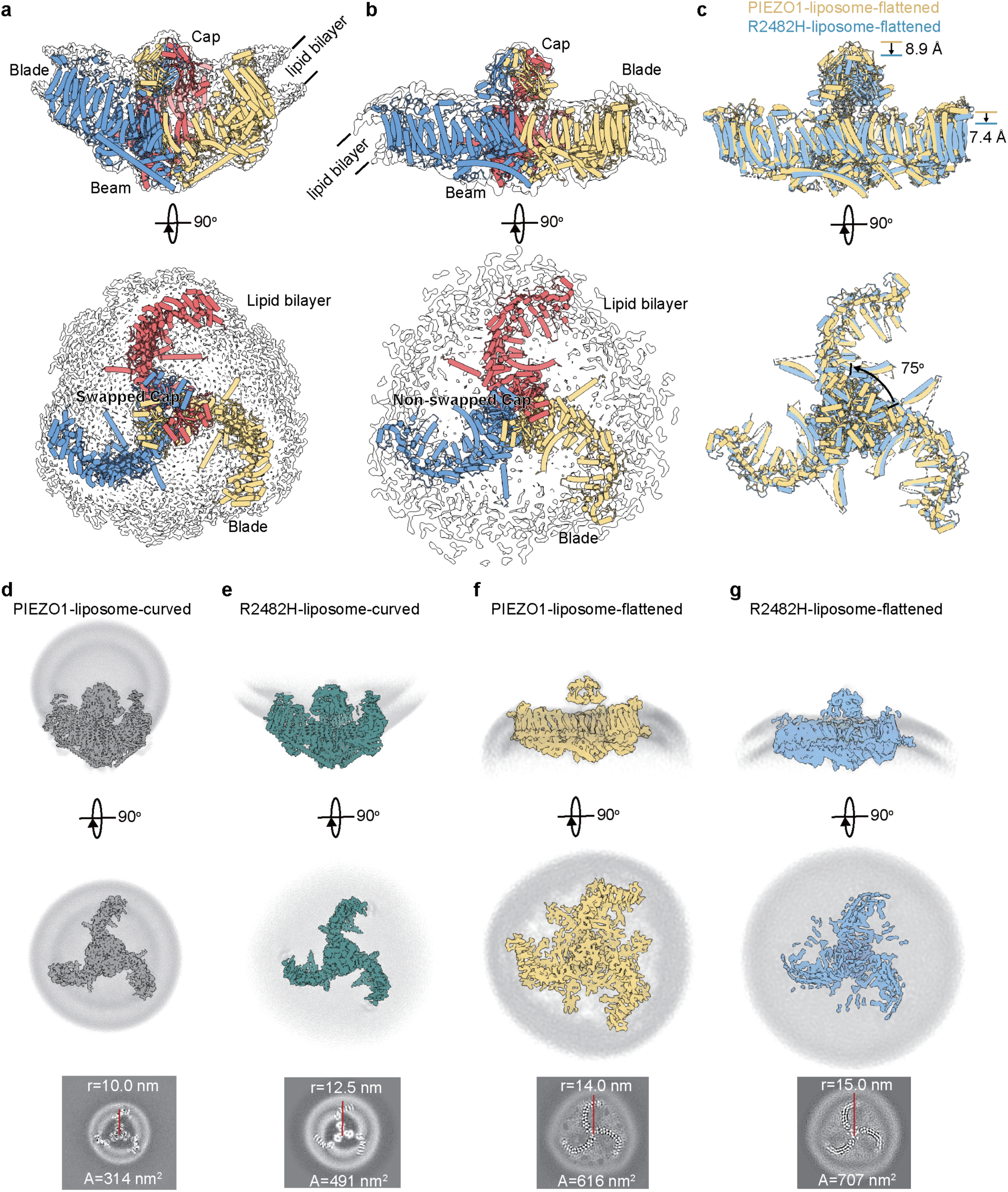
Determination and analyses of R2482H-liposome structures. **a**, Side and top view of the superimposed model with 26 resolved TMs in each subunit and map of the R2482H-liposome-curved structure. Each subunit of the model is shown in different colors. **b**, Side and top view of the superimposed model with 26 resolved TMs in each subunit and map of the R2482H-liposome-flattened structure. Each subunit of the model is shown in different colors. **c**, Side and top view of the superimposed models of the PIEZO1-liposome-flattened (yellow) and R2482H-liposome-flattened (blue). The relative displacement of the cap and blade is labeled. **d**-**g**, The side view (upper row) and projected top view (lower two rows) with the measured projection radius and projection area indicated.

Compared with the previously reported PIEZO1-detergent-curved structures (PDB: 5Z10 and 6B3R) with a measured curvature radius (R) of ∼11.8-12.3 nm, the R2482H-detergent-curved structure is moderately flattened, with an increased R value of 13.7 nm, which is similar to that of the open-prone S2472E-detergent-curved structure (Extended Data Table 1). The difference in the intrinsic curvature suggests that the GOF mutants are more prone to flattening. In line with this reasoning, extensive 3D classification revealed a more flattened structure at a resolution of 6.35 Å, termed R2482H-detergent-partial-flattened (Extended Data Fig. 2), with an increased R value of ∼16.8 nm (Extended Data Table 2). Notably, in the map of R2482H-detergent-partial-flattened, three rod-like densities are laterally located right above the lateral portals (Extended Data Fig. 2 and 6), which likely represent the associated “pore lipids” previously observed in the PIEZO1-liposome-flattened structure^36^ and the PIEZO1-flattened structure from the cell membrane vesicle^37^. Together, these structural analyses suggest that the R2482H mutation induces intrinsic structural changes, including stabilization of the distal blades, and renders the protein more prone to flattening.

### Correlation of the blade height with the constriction of the transmembrane pore

Despite that both PIEZO1 and PIEZO2 form similar homotrimeric architectures^9,10^, they have distinct biophysical properties. Compared to PIEZO1 and PIEZO1-R2482H, PIEZO2 has faster inactivation kinetics and markedly reduced stretch sensitivity^1,38^. The determination of the full structures of both PIEZO2 and PIEZO1-R2482H enabled us to directly compare their structural differences, which might underscore their distinct biophysical properties. Notably, when superimposed on the PIEZO2-detergent-curved structure, the R2482H-detergent-curved structure shows apparent lower but more extended blades. The blade height between the outermost TM3 and the central IHs was measured as ∼9.1 nm and ∼5.8 nm for PIEZO2-detergent-curved and R2482H-detergent-curved structures, respectively (Fig. 1c and Extended Data Table 2). The lack of the distal TM1-12 of THU1-3 in the PIEZO1-detergent-curved structures has precluded accurate measurement of the blade height of PIEZO1. However, based on the observation that the blade height measured at the resolved outmost TM15 of PIEZO1-detergent-curved (∼5.2 nm) is higher than that of R2482H-detergent-curved (∼5.0 nm), but lower than that of PIEZO2-detergent-curved (∼6.6 nm) (Extended Data Table 2), we infer that the distal blade height of PIEZO1 lies between 6 and 9 nm. As shown in the top-down view of the superimposed OH and IH (Fig. 1d), the blade height is strongly correlated with the constriction extent of the IH-enclosed transmembrane pore. As evidenced by the pore radius plot and the CHAP-based hydrophobicity profile of the TM pore of the curved structures of PIEZO1, R2482H, and PIEZO2 (Fig. 1e), L2469 and V2476 in PIEZO1 and the corresponding L2743 and V2750 in PIEZO2 define the hydrophobic TM gate region. PIEZO2 apparently has the most constricted TM pore (Fig. 1e), followed by PIEZO1 and then R2482H. By contrast, the expansion state of the peripheral blades does not show such a strong correlation with the constriction state of the TM pore. For instance, the expansion extent of the blades follows the order of PIEZO1, R2482H and PIEZO2 (Fig. 1c). Notably, the distal end of the full TM-blade of R2482H extends by ∼ 4 nm relative to that of PIEZO2 (Fig. 1c). A recent study has proposed that the modestly flatter blades of the human PIEZO1 (hPIEZO1) relative to those of mouse PIEZO1 (mPIEZO1), might account for the relatively more rapid inactivation kinetics of hPIEZO1^39^. In contradict to the proposal, PIEZO2 has a faster inactivation rate than PIEZO1, but more curved blades. Taken together, these analyses indicate that the flexibility of the blades to be flattened might be a major determinant of the distinct mechanosensitivity between PIEZO1 and PIEZO2, as well as the relatively potentiated mechanosensitivity of R2482H over PIEZO1, but might not have a clear correlation with the inactivation kinetics.

### Curved and flattened structures of R2482H in liposome membranes

The distinct resting curvature states of PIEZO1-curved, S2472E-curved and R2482H-curved prompted us to ask whether the intrinsic curvature property of the PIEZO1 protein itself might dominate its curvature state in lipid membranes. We have previously determined the fully curved PIEZO1-curved-liposome structure by reconstituting PIEZO1 proteins into liposome vesicles made from the soy total lipid extract^36^. Following the same procedure^36^, we reconstituted R2482H into liposomes and determined the structures of the resulting proteoliposomes using cryo-EM. Similar to the PIEZO1-proteolipsome, the majority of the R2482H mutant proteins were preferentially reconstituted into the liposomes in an outside-in configuration, in which the characteristic extracellular cap domain is located inside the lipid vesicles (Fig. 2a,e and Extended Data Fig. 3). By contrast, a minor portion of R2482H-proteoliposomes adopted the outside-out orientation (Fig. 2b,g and Extended Data Fig. 4). In order to obtain a sufficient amount of the outside-out R2482H-proteoliposome vesicles and overcome their conformational instability and heterogeneity, we extensively collected a total of 69,844 micrographs from six datasets. Reconstructing around 272,545 outside-in vesicles led us to determine the R2482H-curved-liposome structure at a resolution of 3.85 Å (Fig. 2a, Extended Data Figs. 3, Extended Data Table 1), which clearly resolved the TM13-TM38, cap and beam domains, 8 “blade lipids” and 1 “pore lipid” in each subunit that are similarly observed in the PIEZO1-liposome-curved structure^36^, as well as the lipid bilayer (Fig. 2e, and Extended Data Fig. 3d). Compared to the nearly spherical shape of the PIEZO1-liposome-curved structure, the R2482H-liposome-curved structure appears flatter, with an increased in-plane radius from ∼10 nm to 12.5 nm (Fig. 2d and e and Extended Data Table 2). Furthermore, while the PIEZO1-liposome-curved structure has 22 resolved TMs (TM17-TM38), the R2482H-liposome-curved structure has 26 resolved TMs (TM13-TM38), consistent with the stabilizing effect of the R2482H mutation on the distal blade. Compared to the R2482H-detergent-curved structure, the R2482H-liposome-curved structure is slightly flattened, resulting in the increase of the R value from ∼13.7 nm to ∼14.7 nm (Extended Data Fig. 5a). Together with the observation that wild-type PIEZO1 also adopts the curvature state in liposome vesicles (R=10 nm) or plasma membrane vesicles^37^ (R=11.8 nm) close to its intrinsic curvature state (R=12 nm), these analyses demonstrate the dominant effect of the intrinsic curvature properties of PIEZO1 and R2482H in controlling their curvature states in lipid membranes.

Multiple rounds of homo and hetero refinement of the small portion of outside-out vesicles picked from the 69,844 micrographs ended up with 65542 vesicles for reconstruction of the R2482H-flatten-liposome map at an overall resolution of 7.45 Å (Extended Data Fig. 4), which is comparable to the previously determined PIEZO1-liposome-flattened structure (6.81 Å)^36^ and the flattened structure of PIEZO1 derived from cell membrane vesicles^37^. The flattened blade domain containing TM13 to TM36 and the levered beam, the anchor domain, and the central pore module were clearly defined, enabling construction of the C_α_ model using the PIEZO1-liposome-flattened model (PDB: 7WLU) as a template^36^. Compared to the PIEZO1-liposome-flattened structure with its cap in the up-state, the cap of the R2482H-liposome-flattened structure is in the down-state, which is similar to the S2472E-intermediate-open structure and the PIEZO1-flatten structure derived from cell membrane vesicles^37^. Remarkably, the OH-Cap-IH pore module of the PIEZO1-liposome-flatten structure shows a ∼75° of counterclockwise rotation in a top-down view (Fig. 2b,c and Fig. 3a). Such a drastic conformational change leads to a switch of the OH-Cap-IH pore module from a swapped organization in PIEZO1 to a non-swapped configuration in the R2482H-liposome-flattened structure (Fig. 2b,c and Fig. 3a). Thus, the R2482H-liposome-flattened structure represents an unexpected and previously unobserved conformational state.

**Fig. 3.**
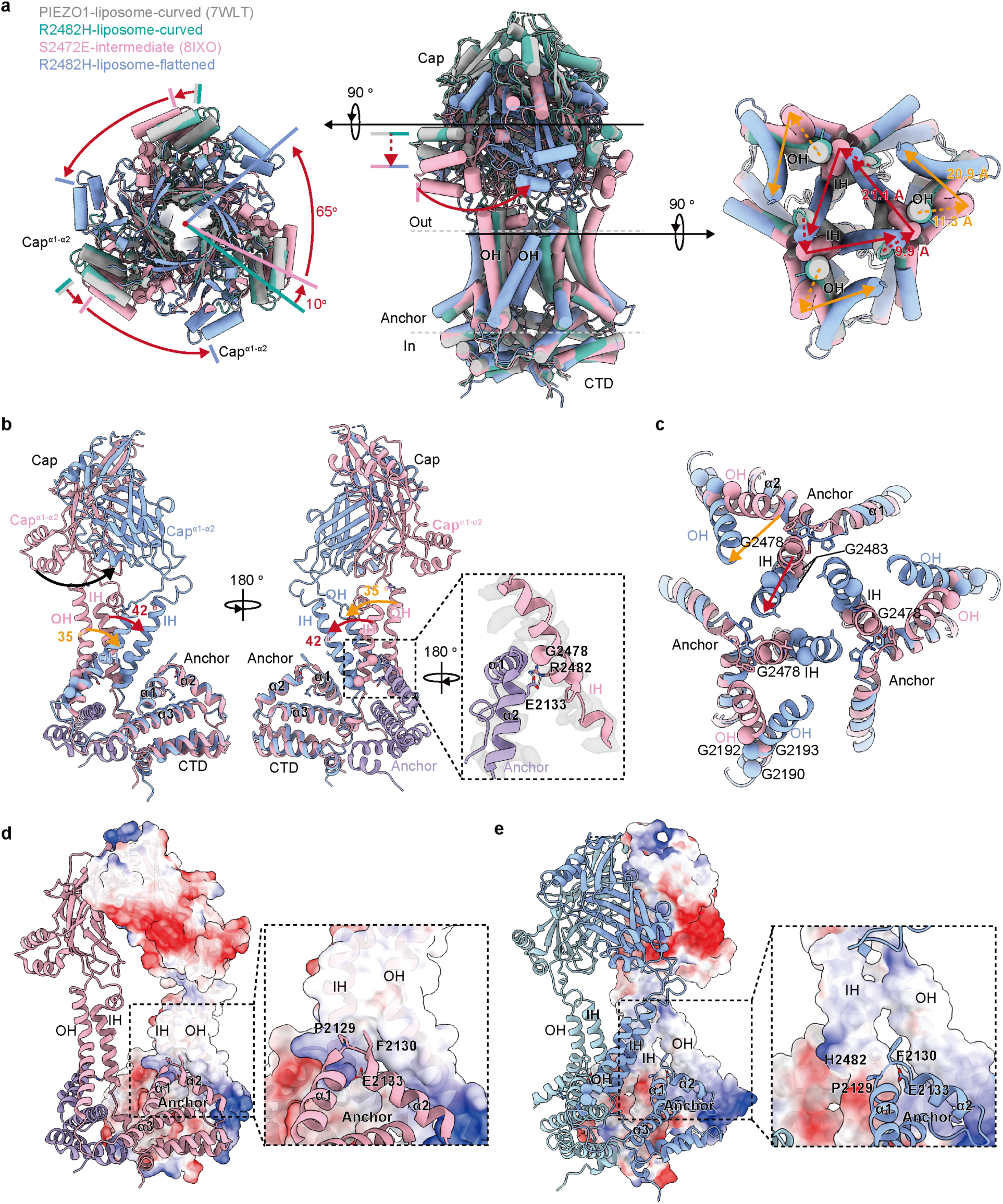
Switching motion of the central pore module. **a**, The top-down views (left and right panels) of the indicated plane and side view (middle panel) of the central pore module of the indicated structures superimposed at the CTD. The dash and solid arrows indicate the step-wise motion of the indicated structural states and the rotational angles of the cap are labeled. **b**, Side view of the Anchor-OH-Cap-IH-CTD domain of the S2472E-intermediate and R2482H-liposome-flattened superimposed at the CTD. The arrows indicate the relative displacement of the Cap^α1-α2^, OH, and IH from the S2472E-intermediate structure to the R2482H-liposome-flattened structure. The dashed box highlights the electrostatic interaction between R2482 in the IH and E2133 in the apex of the swapped anchor domain. **c**, Top-down view showing the indicated displacement of the OH and IH from the S2472E-intermediate structure to the R2482H-liposome-flattened structure. **d**, Side view of the central pore module of the S2472E-intermediate structure showing the interaction among the anchor, OH and IH. The dashed box highlights the closely positioned apex of the anchor domain from one subunit to the packed OH and IH from the neighboring subunit. The highly conserved residues P2129, F2130, and E2133 are labeled. **e**, Side view of the central pore module of the R2482H-liposome-flattened structure showing the less closely associated anchor, OH and IH from the same subunit.

Compared to the PIEZO1-liposome-flattened structure with a R value of ∼109 nm, the blade-beam domain of the R2482H-liposome-flattened structure is further flattened by ∼ 7.4 Å (Fig. 2c), resulting in an increased R value of ∼278 nm (Extended Data Table 2). Thus, the R2482H structure is consistently flatter than the PIEZO1 structure in both curved and flattened conformations. The PIEZO1-flattened structure derived from plasma membrane vesicles was reported to have an additional 2 nm of flattening relative to the PIEZO1-liposome-flattened structure^37^. The difference in lipid compositions between the cell membrane and soy lipid extracts used for liposome generation has been proposed to account for the distinct flatness of the PIEZO1-flattened structures^37^. Based on the observation that stretch-activated currents were only recorded from plasma membrane vesicles derived from PIEZO1-expressing cells, but not from PIEZO1-reconstituted giant unilamellar vesicles made from soy PC lipids, it has been proposed that the observed lipid-like molecules located at the lateral portals might function as a co-factor to enable the mechanical activation of PIEZO1 specifically derived only from cell membranes^37^. However, the observation of similar densities in the PIEZO1-liposome-flattened, R2482H-detergent-partial-flattened, as well as the low-pass map of the R2482H-detergent-curved structures suggests that these densities more likely reflect general lipid association rather than cell-specific cofactors (Extended Data Fig. 6). Additionally, our observation of a flatter R2482H-liposome-flattened structure than PIEZO1-liposome-flattened structure using the exact same soy lipid extracts demonstrates the intrinsic property of PIEZO1 and R2482H proteins themselves in affecting their flatability in lipid membranes. Importantly, the structures of PIEZO1-liposome-flattened, R2482H-liposome-flattened, and PIEZO1-flattened derived from the cell membrane show similar disassembled density of the central plug domain, fragmented density of the lateral plug and latch domain, and fold of the wedge domain (Extended Data Fig. 5b, c)^37^. Therefore, we reason that the extent of the blade flatness of the PIEZO1-liposome-flattened structure might be sufficient to cause unplug of the lateral-plug-gate. However, the lack of side-chain density of the IH in all three flattened structures and different poses of the top cap domain have prevented accurate calculation of the pore radius and a direct comparison of the effect of different degrees of blade flatness on the size and conductivity of the ion-conducting pore.

### Mechanogating parameters of the R2482H mutant channel

On the basis of the PIEZO1-liposome-curved and PIEZO1-liposome-flattened structures (Fig. 2d and f), we have previously estimated the in-plane areal expansion (ΔA) of ∼300 nm^2^ associated with the transition of PIEZO1 from the curved (projected in-plane area of ∼314 nm^2^) to the flattened state (projected in-plane area of ∼616 nm^2^), the force of ∼90 pN, the work of ∼570 pN.nm, the half maximal tension (λ_50_) of ∼1.9 pN.nm^−1^ (Ref^36^). Consistent with being flatter than PIEZO1, the R2482H-liposome-curved and R2482H-liposome-flattened structures have a larger projected in-plane area of ∼491 nm^2^ and ∼707 nm^2^ respectively, generating a projected ΔA of ∼216 nm^2^ (Fig. 2e and g). Given a resting blade height of ∼5.8 nm for R2482H-curved and the previously determined linear regression equation of y = 7.0x + 22 that describes the force-induced displacement of the blade, we deduced a force of ∼60 pN and a work of ∼250 pN.nm for the conformational transition from the R2482H-liposome-curved to the R2482H-liposome-flattened state. Considering the free energy equation ΔG = ΔG (λ=0) -λΔA, where ΔG and ΔG (λ=0) are the free energy difference between the curved and flattened state with or without applied tension (λ). At the equilibrium, where ΔG equals 0, ΔG (λ=0) = λ_50_ΔA = 250 pN.nm, with ΔA is 216 nm^2^. Thus, λ_50_ = 250 pN.nm/216nm^2^ = 1.2 pN.nm^−1^, which is modestly lower than the calculated λ_50_ of PIEZO1 (1.9 pN/nm) and the experimentally measured range of 1.4-4.5 pN/nm^40–42^. The calculated lower λ_50_ of the mutant than that of PIEZO1 is consistent with the patch-clamp-measured modest change of pressure sensitivity^4,5^.

### Effect of R2482H in reorganizing the central pore

The drastic twisting motion of the OH-Cap-IH in the R2482H-liposome-flattened structure led to a reorganization of the central pore from a swapped to a non-swapped configuration. To appreciate the step-wise conformational changes, we superimposed the pore module derived from the structures of PIEZO1-liposome-curved (PDB: 7WLT), R2482H-liposome-curved, S2472E-intermediate-open (PDB: 8IXO), and R2482H-liposome-flatten using the intracellular CTD (D2501-E2547) as the alignment reference (Fig. 3a). The cap of the largely superimposed PIEZO1-liposome-curved and R2482H-liposome-curved shows a 9 Å downward movement and a ∼10° of counterclockwise rotation to reach that of S2472E-intermediate-open, which is further rotated ∼65° to reach that of R2482H-liposome-flattened (Fig. 3a). Concurrently, the upper portion of IH and OH of PIEZO1-liposome-curved and R2482H-liposome-curved display an outward expansion of 9.9 Å and 11.3 Å respectively to reach those of PIEZO1-S2472E-intermediate-open, which then show ∼42 ° and 35° of tilting to reach those of the R2482H-liposome-flattened (Fig. 3a). As a result, the upper end hinge residue G2465 of IH shows a displacement of ∼ 21.1 Å (Fig. 3a). Superimposition of S2472E-intermediate-open and R2482H-liposome-flatten reveals that the IH tilts along its lower end at the flexible residue G2483 right after R2482H, while the OH tilts along its lower portion around the flexible residues G2190, G2192 and G2193 (Fig. 3b,c).

In the PIEZO1-liposome-curved and S2472E-intermediate-open structures, the lower end of IH of PIEZO1 is in proximity to the apex of the anchor domain from the neighboring subunit, which contains highly conserved residues in PIEZO channels, including P2129, F2130 and E2133 (Fig. 3b-d). The positively charged R2482 and the negatively charged E2133 form an electrostatic interaction (Fig. 3b). Mutating E2133 in PIEZO1 reduces the single-channel conductance and the sensitivity to the pore blocker ruthenium red, but slows the inactivation kinetics^43^. The IH leans against the apex of the anchor at the position of G2478, resulting in the side chain benzene group of F2130 fitting right into the hydrophobic pocket enclosed by the IH and OH (Fig. 3b-d). The upper and lower portions of the OH form closely packed hydrophobic interactions with the non-pore-facing side of the IH and the anchor^α2^, respectively (Fig. 3d). Such organization provides stable coupling between IH, OH, and the anchor. The R2482H mutation apparently disrupts the electrostatic interaction with E2133, thereby enabling the drastic tilting motion of the IH at G2483 (Fig. 3b,c). The tilted IH of R2482H-liposome-flattened becomes separated from the apex of the anchor domain and also the upper portion of the tilted OH (Fig. 3e). These structural changes collectively make the IH and OH more mobile and less stably organized, explaining their reduced resolvability in the R2482H-liposome-flattened structure.

### Unaltered ion selection and permeation properties of R2482H

To examine whether the drastic reorganization of the pore module might affect the ion-permeation properties, we characterized the single-channel conductance and ion selectivity of PIEZO1 and R2482H (Extended Data Fig. 7). Our previous studies have revealed that PIEZO1 utilizes the three lateral portals as the intracellular ion permeation routes, which are physically plugged by the three lateral-plug-gates that are strategically latched onto the central axis via the interaction between Y1412 in the latch domain and E2537 in the CTD positioned in the central axis^33^. Such a plug-and-latch mechanism ensures coordinated gating of the three lateral-plug-gates. Mutating Y1412 or E2537 in PIEZO1 disrupted the co-ordinated gating of the three lateral-plug-gates and generated sub-conductance states corresponding to opening of one, two or three of the lateral-plug-gates^33^. To examine whether R2482H-induced reorganization of the pore module might alter the intracellular ion permeation route, we generated R2482H/Y1412A and R2482H/E2537A double mutants and recorded their single-channel currents. Histogram analyses of the single-channel current revealed that PIEZO1 and R2482H exhibit two major current peaks of -3.3-3.4 pA and -2.5 pA and a minor peak of -1.1 pA, which correspond to the 3/3, 2/3 and 1/3 of the full single-channel conductance of PIEZO1 (Extended Data Fig. 7a,b). Compared to PIEZO1 and R2482H, the double mutants R2482H/Y1412A and R2482H/E2537A exhibited a more promenet current peak of -1.1 pA and -0.8 pA respectively, suggesting that Y1412A or E2537A mutation induces similar uncoordinated gating of the three lateral-plug-gates in both PIEZO1 and R2482H (Extended Data Fig. 7a,b). Thus, like PIEZO1, R2482H similarly utilize the three lateral portals and lateral-plug-gates as the intracellular ion permeation routes. We further recorded the single-channel currents of PIEZO1 and R2482H at different voltages and found that they had similar slope single-channel conductances, suggesting that R2482H does not affect the ion permeation property (Extended Data Fig. 7c,d).

Combining the whole-cell patch clamp electrophysiology and the piezo-driven indentation of the cell membrane with a blunt glass pipette, we measured the mechanically activated currents under varied voltages in the designed ionic condition for measuring the permeability ratio of Cl^−^ to Na^+^ (P_Cl_/P_Na_) (Extended Data Fig. 7e, f). On the basis of the current-voltage relationship and the calculated reversal potential, we obtained similar P_Cl_/P_Na_ for PIEZO1 and R2482H, suggesting that R2482H does not affect the ion selectivity (Extended Data Fig. 7e, f).

These electrophysiological characterizations collectively demonstrate that the ion permeation and selection properties of P2482H are not altered by the reorganization of the pore modue from a swapped to a non-swapped configurations.

### Conformational transitions of R2482H from the flattened-twisted to the curved-partial-twisted state

A recent study attempted to determine the structures of several DHS-associated human PIEZO1 mutants, including hPIEZO1-R2456H (the corresponding mutant of mPIEZO1-R2482H)^39^. In contrast to our present study, the hPIEZO1-R2456H protein itself appeared to be unstable for structural determination under their experimental conditions. However, a medium-resolution structure of hPIEZO1-R2456H in complex with the MyoD-family inhibitor protein MDFIC was determined. When co-expressed with PIEZO1, MDFIC drastically slowed the inactivation and nearly abolished the deactivation of the stretch-induced currents^44^. Notably, when superimposed with the R2482H-liposome-flattened and R2482H-liposome-curved structures, the hPIEZO1-R2456H/MDFIC-curved structure (PDB: 8YFG) shows curved blades (Fig. 4a), while the OH-Cap-IH pore module is aligned in between (Fig. 4b). We interpretate the hPIEZO1-R2456H/MDFIC-curved structure might represent a transition state from the R2482H-liposome-flattened to the R2482H-liposome-curved, in which the blades transit from the flattened state back to the curved state, while the twisted OH-Cap-IH pore module transits from the non-swapped position back to the swapped configuration. We term the hPIEZO1-R2456H/MDFIC-curved structure R2456H/MDFIC-curved-partial-twisted. Given that MDFIC rendered the PIEZO1-mediated current nearly non-deactivating upon removal of mechanical stimuli^44^, we reason that the R2456H/MDFIC-curved-partial-twisted structure might represent a transition state in the process of deactivation, in which the flattened blade returns to the curved state, while the OH-Cap-IH pore module switches from the non-swapped configuration back to the swapped configuration.

**Fig. 4.**
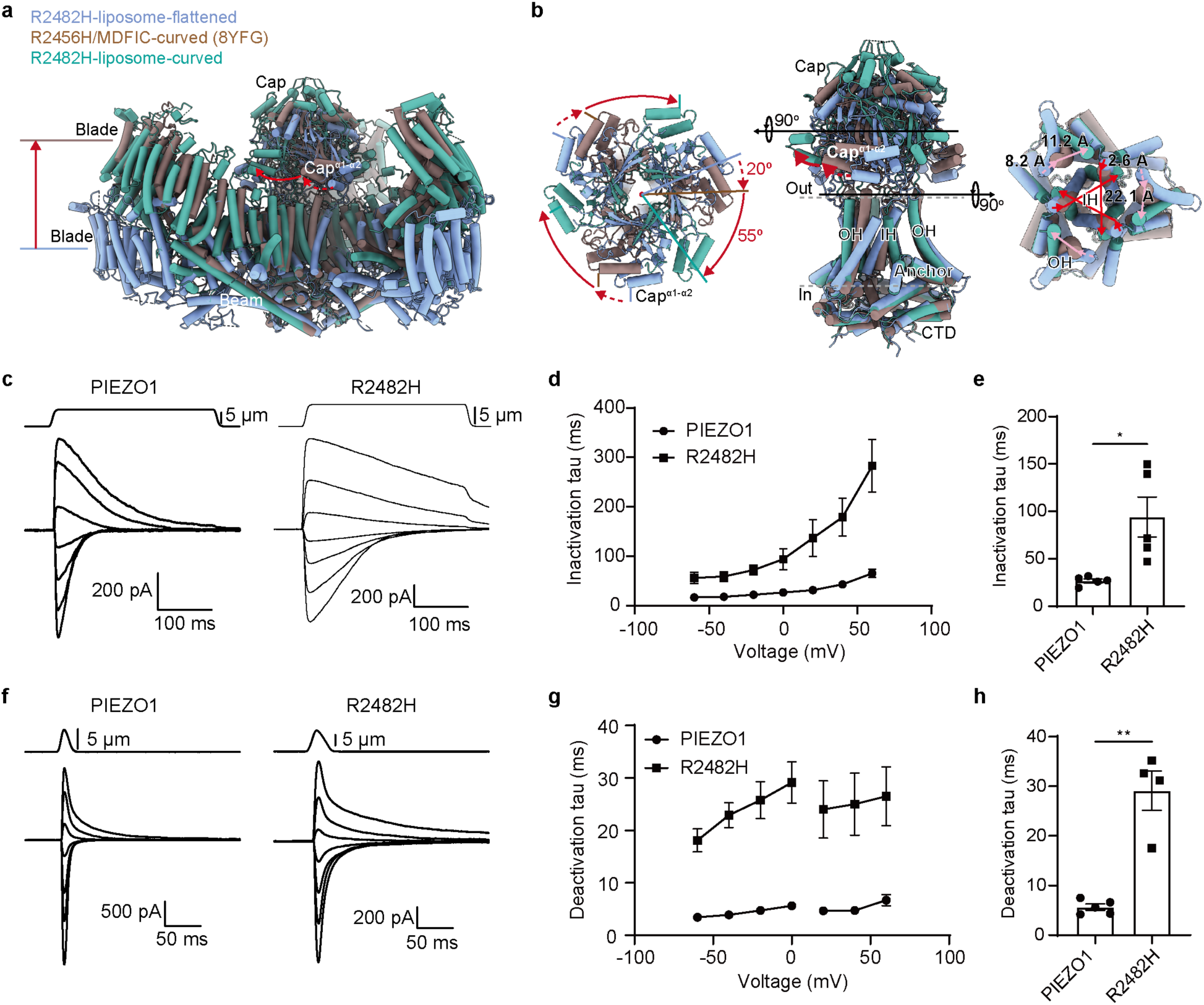
Deactivation transitions and gating kinetics of R2482H. **a**, Side view of the indicated structures superimposed at the CTD. The dash and solid arrows indicate the step-wise motion of the indicated domains. **b**, The top-down views (left and right panels) of the indicated plane and side view (middle panel) of the central pore module of the indicated structures superimposed at the CTD. The dash and solid arrows indicate the step-wise motion of the indicated domains. **c**, Representative traces of whole-cell currents from cells transfected with PIEZO1 or R2482H in response to 300 ms of poking stimulation under varied voltages ranging from -60 mV to 60 mV in 20 mV increments. **d**, Averaged inactivation tau of PIEZO1- or R2482H-mediated poking-evoked currents recorded at the indicated voltages. **e**, Scatter plots of the inactivation tau of PIEZO1- or R2482H-mediated poking-evoked currents recorded at 0 mV. Each bar represents mean ± SEM (n=4-5 cells). Unpaired Student’s t-test with Welch’s correction, *p < 0.05. **f**, Representative traces of whole-cell currents from cells transfected with PIEZO1 or R2482H in response to a brief poking stimulation under varied voltages ranging from -60 mV to 60 mV in 20 mV increments. **g**, Averaged deactivation tau of PIEZO1- or R2482H-mediated poking-evoked currents recorded at the indicated voltages. **h**, Scatter plots of the deactivation tau of PIEZO1- or R2482H-mediated poking-evoked currents recorded at 0 mV. Each bar represents mean ± SEM (n=4-5 cells). Unpaired t-test with Welch’s correction, **p < 0.01.

### Slowed inactivation and deactivation kinetics of R2482H

To functionally support the proposed structural dynamics of deactivation, we compared the inactivation and deactivation kinetics of PIEZO1- and R2482H-mediated whole-cell currents recorded under varied voltages from -60 mv to 60 mV. As shown in Fig. 4c, to characterize the inactivation process, we applied 300ms of poking stimulation and recorded the mechanically activated currents with inactivation. Compared to PIEZO1, R2482H exhibited apparently slowed inactivation kinetics and increased inactivation Tau values across all the tested voltages (Fig. 4c-e). For example, at 0 mV, the inactivation Tau of R2482H (94.0 ± 21.1 ms) is significantly larger than that of PIEZO1 (26.8 ± 2.1 ms) (Fig. 4e). To characterize the deactivation process, a poking stimulation was applied at a speed of 1 μm/ms and immediately withdrawn once a robust current was elicited. The mechanically activated currents and the decay of the current were subsequently recorded. (Fig. 4f). Compared to PIEZO1, R2482H showed markedly slowed deactivation kinetics and increased deactivation Tau values across all the tested voltages (Fig. 4f-h). For example, at 0 mV, the deactivation Tau of R2482H (29.1 ± 4.0 ms) is significantly larger than that of PIEZO1 (5.6 ± 0.6 ms) (Fig. 4h). These electrophysiological characterizations demonstrate that the inactivation and deactivation kinetics of R2482H is markedly slower than that of the wild-type PIEZO1. At 0 mV under which the structure was determined, a portion of the R2482H-mediated current remained in either the inactivation or the deactivation processe. This might explain the capture of the R2482H-flattened-twisted structure and the deactivation-transition state represented by the hPIEZO1-R2456H/MDFIC-curved-partial-twisted structure (Fig. 4a).

### Distinct conformational transitions between PIEZO1 and R2482H

On the basis of the varied conformational states of PIEZO1-curved (R of ∼10-12 nm), S2472E-curved or R2482H-curved (R of ∼14 nm), R2482H-partial-flattened (R of ∼17 nm), S2472E-intermediate-open (R of 32 nm), PIEZO1-liposome-flattened (R of 109 nm), R2482H-liposome-flattened-twisted (R of 278 nm) (Extended Data Table 2), R2456H/MDFIC-curved-partial-twisted, we deduced the stepwise conformational transitions of PIEZO1 and R2482H (Fig. 5a and Supplementary Videos 1-4). The structural dynamics of PIEZO1 starts from the intrinsically curved state represented by the PIEZO1-curved structure, which migh have a flexibility to adopt a range of curvature radius in the lipid membranes^34,36,45–47^. The downward movements of the cap, together with the partial flattening of the blades, lead to an intermediate state represented by the PIEZO1-S2472E-intermediate-open structure, in which the cap gate and TM gate are open, while the intracellular lateral plugs remain plugged. A further flattening of the blades and lever-like bending and sliding of the beams of the S2472E-intermediate-open structure is expected to transit to the state represented by the combination of the central pore module of the PIEZO1-S2472E-intermediate-open structure and the flattened blade-beam domain of the PIEZO1-flatten-liposome structure, in which the cap-gate, TM-gate and intracellular lateral plug-gate are expected to open (Supplementary Videos 1-2). In line with this prediction, the reported PIEZO1-flattened structure derived from cell membranes likely represents the presumed open state that has fully flattened blades (2 nm of additional flattening of the blades relative to the PIEZO1-liposome-flatten structure) and down-state cap^37^. Such a conformational state might readily transit to the state represented by the PIEZO1-liposome-flattened structure, whose blades remain flattened, while the cap is released back to the up-state via the spring-like motion of the OH-cap-linkers and Cap-IH-linkers that connect the cap to the OH and IH, respectively, resulting in channel inactivation (Supplementary Videos 1-2). Upon removal of mechanical stimuli, during deactivation, the blades return from the flattened to the curved state, while the cap remains in the up-date, resulting structural transition back to the curved-closed state of PIEZO1 (Supplementary Videos 1-2).

**Fig. 5.**
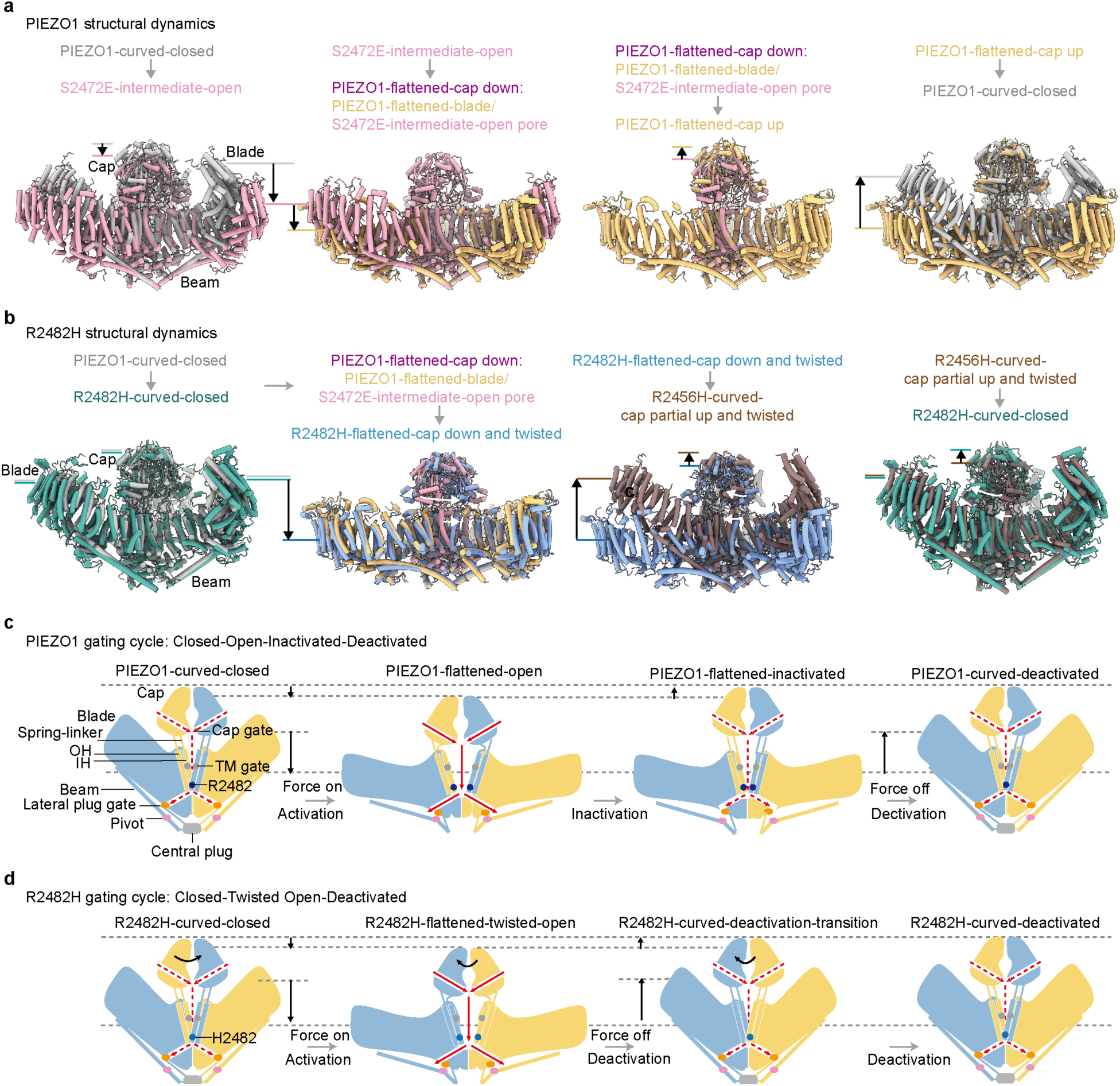
Step-wise conformational transitions of PIEZO1 and R2482H. **a** and **b**, The structural transition cycle of PIEZO1 (**a**) or R2482H (**b**) between the curved and flattened states. The gray arrows indicate the transitional state, while the black and white arrows indicate the relative displacement of the indicated domains. **c** and **d**, The proposed gating cycles of PIEZO1 (**c**) and R2482H (**d**) are illustrated in two out of the three subunits. Due to the medium resolution of the structures representing the PIEZO1-flattened-open, PIEZO1-flattened-inactivated, and R2482H-flattened-twisted open, the pore size of the presumed conducting pore remains ambiguously determined.

Compared to PIEZO1-curved, the intrinsically flatter R2482H-curved (as well as the S2472E-curved) structure is more predisposed to flattening, accounting for its potentiated mechanosensitivity. Like PIEZO1, the R2482H mutant might transit through the R2482H-partial-flattened state, the intermediate state represented by the S2472E-intermediate-open, and then the fully flattened state with cap in the down-state (Fig. 5b). An additional twisting motion of the central pore module leads to the state represented by the R2482H-liposome-flattened-twisted structure, in which the central pore switches from the swapped to the non-swapped configuration (Supplementary Videos 3-4). The reorganization of the central pore region might disrupt the spring-like transition of the cap from the down-state to the up-state, thus preventing the transition to the inactivated state represented by the PIEZO1-liposome-flattened, in which the blades remain flattened, while the cap is in the up-state for closing the cap-gate. Instead, upon removal of mechanical stimuli, the R2482H-liposome-flattened structure transits to the state represented by the R2456H/MDFIC-curved-partial-twisted state, in which the blades recurve, while the central pore module is in the middle of twisting back from the non-swapped to the swapped configuration. A fully twisting and upward motion of the pore module back to the swapped configuration results in the completed conformational changes to the R2482H-curved state (Supplementary Videos 3-4).

## Discussion

The mechanically activated PIEZO channels exemplify the intricate relationship between protein structure, gating dynamics, and physiological function. Mutations that alter gating kinetics, such as the DHS-associated R2482H in PIEZO1, can lead to profound pathophysiological consequences, including red blood cell dehydration and iron overload. In this study, our structural determinations and analyses reveal that a single residue mutation of R2482H out of 2547 residues of PIEZO1 can lead to a drastic change in the overall channel structure, including the long-distance stabilization of the distal blade and the reorganization of the central pore module from a swapped to a non-swapped configuration. Similar point mutation-induced drastic changes in PIEZO1 structure have also been observed in the open-prone S2472E mutant, which stabilizes an intermediate open conformation. These examples might highlight the identification of many missense mutations in PIEZO1 in association with human genetic diseases^48,49^. Combining the strength of mutational effects and membrane vesicle-derived forces on PIEZO1, we have determined a series of conformational states of PIEZO1, which consequently allows us to deduce the structural dynamics underlying the complete gating cycle of activation, inactivation and deactivation, as well as to understand the altered structure-function relationship of the disease-causing mutations such as R2482H (Fig. 5 and Supplementary Videos 1-4).

A central finding of our work is the determination of the complete 38-TM-topology structure of PIEZO1, achieved through the R2482H-detergent-curved state (Fig. 1). The finding that a single point mutation R2482H at the pore-lining IH can allosterically stabilize the highly flexible distal TM1-TM12 regions, located ∼25 nm away, underscores the remarkable conformational coupling across the entire protein. This suggests that the N-terminal domain, while unresolved in structures of wild-type PIEZO1, is an integral component of the blade and is likely involved in transmitting mechanical force. In line with this, we have found that a fetal hydrops-associated L322P mutation in the TM8 of hPIEZO1 (or L329P in mPIEZO1) abolished the mechanosensitivity of PIEZO1 while retaining activation by the chemical activators Yoda1 and Jedi1^50^. Deleting the extracellular loop 15-16 or 19-20 or mutating the pivot residues L1342 and L1345 in the beam also affects the mechanical activation of PIEZO1^51^. These studies suggest a reciprocal mechanical transduction between the peripheral blade and the central pore. Notably, the intrinsic curvature (R=14-15 nm) of the R2482H mutant is flatter than wild-type PIEZO1 (R=10-12 nm) in both detergents and lipid membranes, indicating the dominant effect of the intrinsic elastic property of PIEZO1 and R2482H proteins in determining their curvature states. The pre-flattened resting state of GOF mutants such as R2482H and S2472E provides a straightforward structural explanation for the mutant’s potentiated mechanosensitivity, as it would require less energy to reach more flattened, open conformations. Indeed, we find a strong correlation between the blade height and the constriction of the transmembrane pore. By comparing the full structures of PIEZO2 and PIEZO1-R2482H, we found that a higher blade curvature correlates with a more constricted pore (Fig. 1c-e). This observation positions the overall blade curvature as a critical determinant of the TM gate’s status, linking the global conformational change of the nano-bowl to the local control of the ion conduction pathway. Indeed, the calculated ΔG_0_ of 250 pN.nm and λ_50_ of 1.2 pN.nm^−1^ for R2482H is quantitatively lower than those for PIEZO1 (570 pN.nm and 1.9 pN.nm^−1^, respectively).

The most striking conformational state we captured is the R2482H-liposome-flattened structure, which exhibits a drastic ∼75° twisting of the central pore module, switching it from the canonical swapped configuration to a novel non-swapped configuration (Fig. 2 and Supplementary Videos 3-4). This reorganization is facilitated by the disruption of a key electrostatic interaction between R2482 and E2133 on the anchor domain of the neighboring subunit. The loss of this “molecular clutch” renders the inner and outer helices more mobile, permitting large-scale twisting motion. A recent study of the structures of hPIEZO1 and hPIEZO1-R2456H/MDFIC has proposed that a “pore lipid” molecule within the central cavity directly interacts with R2456, forming a lipid-phosphate-arginine contact that stabilizes the closed state, whereas the R2456H mutation disrupts this interaction, resulting in displacement of the IH^39^. However, given that no apparent lipid density was observed in the reported medium-resolution map of the R2456H/MDFIC complex and the existence of lipid-like density also in our R2482H structures (Extended Data Fig. 6), the relevance of lipids in functionally regulating the inactivation kinetics of PIEZO1 and hPIEZO1-R2456H or mPIEZO1-R2482H and structurally remodeling the central pore module remains illusive. Regardless the involvement of lipids, on the basis of the slowed inactivation kinetics (Fig. 4f-g), the association of the down-stated cap with the opening of the cap gate, the apparent expansion of the upper portion of the IHs, disassembly of the central plug, fragmentation of the lateral-plug-gate, and folding of the wedge domains, we propose that the R2482H-liposome-flattened-twisted structure might likely represent a distinct open state from the presumed open state of PIEZO1 characterized by flattened blades and down-stated cap, which might be represented by the PIEZO1-flattened structure derived from cell membrane vesicles (Fig. 5c, d). Nevertheless, the lack of side chain density of the two presumed open structures has prevented us from precisely measuring the pore radius and ion permeation status.

On the basis of the distinct conformational states of PIEZO1 and R2482H, we propose distinct structural dynamics underlying their distinct gating properties (Fig. 4c-g, Fig. 5c, d, and Supplementary Videos 1-4). For wild-type PIEZO1 (Fig. 5c and Supplementary Videos 1-2), the PIEZO1-curved-closed state can be mechanically induced into the PIEZO1-flattened-open state by lowering the cap, flattening its blades, levering the beams, dissembling the central plug, leading to the opening of the cap-gate, TM-gate and lateral plug-gate. The open state rapidly transitions to the inactivated state by just releasing the cap from the down-state back to the up-state, closing the cap gate for inactivation. Combining comprehensive mutagenesis and electrophysiological characterizations, we have shown that the OH-Cap-linkers and Cap-IH-linkers function as entropic springs to drive activation, inactivation, and stochastic single-channel gating of PIEZO1. Upon removal of mechanical stimuli, the channel is deactivated by returning the blade from the flattened to the curved state, resulting in the complete structure-function dynamic cycle.

As for the R2482H mutant (Fig. 5d and Supplementary Videos 3-4), the intrinsically flatter R2482H-curved structure can be more readily induced by mechanical forces to the R2482H-flattened-twisted-open state. The reorganization of the pore module into a non-swapped configuration likely disrupts the spring-like mechanism that, in wild-type PIEZO1, allows the cap to return to the up-state for rapid inactivation. Instead, the mutant channel becomes trapped in the R2482H-flattened-twisted-open state, unable to rapidly transition to an inactivated state, explaining its slowed inactivation kinetics. Upon removal of mechanical forces for deactivation to occur, the R2482H-flattened-twisted-open state might go through an R2482H-curved-deactivated-transition state represented by the R2456H/MDFIC-curved-partial-twisted structure, where the blades have re-curved, but the pore module remains partially twisted and has not fully returned to the swapped configuration, resulting in slowed deactivation kinetics. Such a structural mechanism aligns with the nearly abolished deactivation kinetics observed when MDFIC is co-expressed with PIEZO1^44^. For the R2482H mutant alone, the slowed deactivation can be explained by a similar, albeit less pronounced, trapping in this transitional pathway.

In summary, our study on the PIEZO1-R2482H mutant has provided profound insights in understanding the structure-function relationship and disease-causing mechanism of PIEZO1. We have revealed the complete topological architecture of PIEZO1, and discovered a novel non-swapped pore conformation that highlights the dynamic plasticity of the central pore module, and elucidated the structural basis for its slowed inactivation and deactivation kinetics. These findings significantly advance our understanding of the gating cycle of PIEZO1, demonstrating how allosteric coupling across vast distances and dramatic conformational changes in the pore work in concert to translate membrane mechanics into precisely controlled ionic signals.

## Acknowledgments

We thank the Protein Preparation and Identification Facility at Technology Center for Protein Science in Tsinghua University for facility support. This work was supported by grant numbers 2021ZD0203301, 32425003, 32130049, 32021002, 31825014 to B.X. from either the National Key R&D Program of China or the National Natural Science Foundation of China, the New Cornerstone Investigator Program, and the Beijing Outstanding Young Scientist Program grant to B.X.

## Author contributions

C.W. and Z.Z. determined the R2482H-liposome structures; X.Y. and S.L determined the single-particle R2482H structures; J.Y. carried out electrophysiological studies; C.L helped cryo-EM studies; X.Z. and B.T. helped with analysis of the structural model; X.L. supervised cryo-EM data collection and image processing; B.X. conceived and directed the study, obtained funding, analyzed the structure, and wrote the manuscript with help from all the authors.

## Competing interests

Authors declare that they have no competing interests.

## Data and materials availability

The structural coordinates of the R2482H-less-Curved, R2482H-detergent-curved, R2482H-detergent-partial-flatten, R2482H-liposome-curved, R2482H-liposome-flattened structures were deposited in the Protein Data Bank under the ID xxxx, xxxx, xxxx and xxxx, respectively. The corresponding cryo-EM maps were deposited into the Electron Microscopy Data Bank (EMDB) under the accession ID EMD-xxxxx, EMD-xxxxx, EMD-xxxxx, and EMD-xxxxx, respectively.

## Methods

### Cell lines

HEK293T cells were used for protein expression and purification. These cells were cultivated in Dulbecco’s Modified Eagle Medium (DMEM) supplemented with 10% fetal bovine serum (FBS), 100 U/ml penicillin, and 100 μg/ml streptomycin. For electrophysiology experiments, we utilized Piezo1-KO-HEK293 cells, where the endogenous Piezo1 gene was disrupted^13^, obtained from Dr. Ardem Patapoutian’s or Dr. Jie Xu’s laboratory. These cells were also cultured in DMEM supplemented with 10% FBS, 100 U/ml penicillin, and 100 μg/ml streptomycin. Transfected cells were briefly digested with a 20-time diluted 0.05% trypsin in 1 ◻ DPBS and reseeded onto poly-D-lysine-coated coverslip 2 (for recording poking-evoked maximal currents) or 12 (for cell-attached configuration) hours before recording.

### Molecular Cloning

For protein purification, the mouse PIEZO1-R2482H encoding sequences were fused with a C-terminal glutathione S-transferase (GST) tag with a precision protease cleavage site in between, followed by the GFP-coding sequence driven by an internal ribosome entry site (IRES) (for monitoring the efficiency of transient transfection and protein expression). For electrophysiology recordings, the PIEZO1, PIEZO1-R2482H, PIEZO1-R2482H/Y1412A, PIEZO1-R2482H/E2537A were fused directly with the mRuby2 fluorescent protein-encoding sequence to accurately reflect the expression and subcellular location of the PIEZO1 proteins.

### Expression and purification of the R2482H mutant proteins

PIEZO1-R2482H was purified following published procedures^30,36^ with two modifications: L-α-phosphatidylcholine was replaced by Soy Total Lipid Extract during membrane solubilization, and the detergent composition was adjusted. Briefly, 36 h after transfecting HEK293T cells with the R2482H-pp-GST-IRES-GFP construct encoding a PIEZO1 R2482H–GST fusion with a PreScission protease site, cells were harvested and lysed for 2 h in 25 mM Na-PIPES (pH 7.2), 150 mM NaCl, 3 mM DTT, 1% (w/v) CHAPS, 0.1% C12E10, 0.05% GDN, 0.5% Soy Total Lipid Extract, and protease inhibitors (Roche). The lysate was clarified (14,000 × g, 30 min), and the supernatant was incubated with glutathione–Sepharose at 4 °C for 6 h. Bound protein was released by on-bead PreScission protease cleavage (0.1 mg ml⁻¹, overnight at 4 °C) and further purified by size-exclusion chromatography (Superose 6 Increase 10/300 GL, GE Healthcare) in 25 mM Na-PIPES (pH 7.2), 150 mM NaCl, and 0.04% C12E10. Fractions eluting at 10.75– 11.75 ml, corresponding to well-dispersed trimeric R2482H (Extended Data Fig. 1a), were pooled for cryo-sample preparation.

### Reconstitution of R2482H proteoliposomes

R2482H was reconstituted into liposomes following the previously described procedure^36^. 40 mg Soy Total Lipid Extract (Avanti) was dissolved in chloroform and dried in a rotary vacuum evaporator at 37 °C for 4h to remove residual chloroform. The soy lipid film was resuspended with 1 ml reconstitution buffer (25 mM PIPES-Na, pH 7.4, 150 mM NaCl) by 3 freeze-thaw cycles using liquid nitrogen and a 40 °C water bath. The lipid solution was then subjected to another 8 freeze-thaw cycles, followed by 31 repeated extrusions through 100 nm filters (Nucleopore) using an Avanti Mini-Extruder. The extruded liposomes were flash-frozen in liquid nitrogen for stock. For reconstitution, the liposome solution made from 3.2 mg soy lipid was solubilized with 2% decyl maltoside (DM) and incubated with 0.8 mg purified R2482H protein at 4 °C on a rotator for 1 h. The protein-lipid-detergent mixture was subsequently loaded on a gravity column packed with 20 mL Sephadex G-50 Superfine resin (Cytiva), and the fractions containing reconstituted proteoliposomes were collected.

The resulting proteoliposomes containing empty liposomes can be further separated by discontinuous density gradient centrifugation as previously described^36^. The collected fractions from the G-50 column were pooled and concentrated to about 250 μL (Amicon Ultra centrifugal filter, 4 ml, 100-kDa cut-off). The discontinuous gradient was constructed by iodixanol (Sigma-Aldrich) (30%, 20%, 15%, 5%, 3%, 1%, 0% from bottom to top). The reconstituted sample was set at the 20% layer and ultracentrifuged at 50,000 g at 4 °C for 16 h in a TLS55 rotor. After ultracentrifugation, the empty liposomes floated into the layers of 3-5%, and the proteoliposomes were mainly condensed at the layers of 15-20%. The 20% layer was collected and the iodixanol was removed by 7 repetitive dilution– concentration cycles (1:5 dilution). The sample was finally concentrated to about 20 μl (Amicon Ultra centrifugal filter, 0.5 ml, 100-kDa cut-off) for cryo-EM sample preparation.

### Sample preparation and cryo-electron microscopy data acquisition

For R2482H in detergent, 4 μl of freshly purified Piezo1-R2482H proteins (approximately 3-41 mg ml-1) each time were sequentially blotted three times to 300-mesh holey carbon Au R1.2/1.3 grids treated with 25s of glow discharge (Quantifoil, Micro Tools GmbH). The grids were then blotted and plunged into liquid ethane using FEI Mark IV Vitrobot operated at 8°C and 100% humidity.

For R2482H proteoliposome, the Quantifoil Au grids (Holey gold, R1.2/1.3, 200mesh) were pre-treated with a prolonged glow discharge time for 40 s. 4 μl freshly prepared R2482H proteoliposomes were loaded onto the grid three times. Each time, wait for 2 minutes, and add the Fluorinated Fos-choline-8 (0.65 mM; Hampton) to the sample before the final loading. Both samples were then blotted for 5 s and plunged into liquid ethane using an FEI Mark IV Vitrobot operated at 8 °C and 100% humidity.

The grids were transferred to a 300 kV Titan Krios (FEI) electron microscope equipped with a GIF Quantum energy filter (slit width 20 eV). Images were recorded using a K3 Summit direct electron detector (Gatan) in super-resolution mode. Data acquisition was performed using AutoEMation II at a nominal magnification of 64,000×, resulting in a super-resolution pixel size of 0.549 Å on the image plane, with defocus ranging from -1.5 to -2.0 μm. The total exposure time was set to 2.56 s with 0.08 s per frame, yielding a 32-frame stack. The total dose was approximately 50 e^−^/Å^2^ for each micrograph.

### Image processing of R2482H in detergent

A simplified flowchart of the procedure for image processing is presented in Extended Data Fig. 2. In the initial stages, three sets of cryo-EM data were processed, comprising 10,572, 6,945 and 10,658 micrograph stacks. The stacks were motion-corrected and dose-weighted using MotionCor2^52^ with 2 × 2 binning, resulting in a pixel size of 1.098 Å. After whole-image CTF estimation using CTFFIND4.1^53^, 9,092, 6,022 and 9,156 good micrographs were manually selected. After auto-picking using RELION-3.1^54^ and several rounds of 2D classification, a total of 493,759, 1,384,800 and 2,131,100 particles were selected and subjected to 3D refinement, resulting in three electron density maps at 7.83 Å, 5.45 Å and 4.53 Å resolution. Three sets of particles were joined for multi-reference 3D classifications performed with four classes. Particles from the highest-resolution class were subjected to 3D refinement, resulting in a density map at 4.18 Å. The PIEZO1 electron microscopy map (EMD-6865), low-pass-filtered to 60 Å, was used as the initial model. Each particle was re-centered and its local defocus parameters were refined using Gctf^55^. The particles that were well-centered and had more accurate defocus parameters were re-extracted from the motion-corrected and dose-weighted micrographs. All 3D refinement and 3D classification were performed with C3 symmetry.

Focused refinements were performed as the distal blades were absent in the overall 4.18 Å map. The subroutine of relion_particle_symmetry_expand in RELION-3.1 was used to expand 342,328 particles with C3 symmetry. Then, 1,026,984 blade particles were obtained using the subtraction of two adjacent blades in raw images. These blade particles were subjected to 3D refinement, resulting in the final focused map of the distal blade at 3.60 Å resolution. As the density of THU1-3 was still fragmentary and unclear in this map, more surrounding regions of the distal blade were subtracted from the 1,026,984 blade particles and subjected to local angular search 3D refinement, yielding the final focused map of the distal blade at 4.07 Å resolution.

As the cap was fragmentary in the map, the focused refinement of the central region of the 342,328 particles was performed by subtracting the projections of most parts of the three surrounding blades. C3 symmetry was imposed during the refinement with a local mask and resulted in a focused map at 3.48 Å resolution. The two final focused maps were fit together according to their overlapping area using Chimera^56^ and then combined using PHENIX^57^ combine_focused_maps. The final density map was created by combining three copies of the combined map.

The reported resolutions are based on the gold-standard Fourier shell correlation 0.143 criterion^58^. All density maps were sharpened by applying a negative B-factor that was estimated using automated procedures^59^. Variations in the local resolution were estimated using Resmap^60^.

### Image processing of R2482H proteoliposome

Simplified schemes of the procedure for image processing are presented in Extended Data Fig. 3,4. A total of 34,579 and 35,265 cryo-EM micrographs were collected from the samples using Quantifoil grids. All the micrographs were motion-corrected using the MotionCor2 program with an 11 × 11 patch and a two-fold binning, resulting in a pixel size of 1.098 Å. Contrast transfer function (CTF) estimation and particle picking were performed by CryoSPARC. First, about 231 and 344 particles in curved state and the flattened state were picked by Manual Picker and subjected to 2D classification, which was used to generate templates for further particle picking in more data. For all the picked particles, visually good ones were further selected and re-centred to the PIEZO1 complexes by analyzing the image alignment results from 2D classification. Finally, 1,109,490 and 247,377 particles in the curved state and the flattened state, respectively, were selected for further 3D analysis.

The sub-dataset of the PIEZO1-R2482H in the curved state was processed as follows. A PIEZO1 map (EMD-6865) was low-pass filtered to 30 Å and used as the initial model. Then, a map reported at 7.39 Å resolution was calculated and revealed the cap domain and the bottom of the beam and transmembrane gate domain. Then, iterative heterogeneous refinement and homogeneous refinement were carried out. In each heterogeneous refinement, those classes with poor PIEZO1-R2482H density were removed, and the best class was used as the initial model of the next round. Finally, 272,545 particles remained and the resolution was pushed to 3.85 Å by non-uniform refinement and local refinement.

The data processing for the PIEZO1-R2482H in the flattened state was similar to that in the curved state. First 132,164 particles in flattened state selected by 2D classification were used to generate the initial model by ab-initio reconstruction. Then iterative homogeneous refinement and heterogeneous refinement were used to remove poor particles. Finally, 65,542 particles were retained in non-uniform refinement to produce a map with an overall resolution of 7.45 Å, revealing the cap domain and the bottom of the beam and transmembrane gate domain.

All the 3D refinement and 3D classification were performed with C3 symmetry. The reported resolutions were based on the gold-standard Fourier shell correlation at 0.143 criterions.

### Model building and structure refinement

For the R2482H-detegent-curved, R2482H-liposome-curved and R2482H-liposome-flattened, the determined cryo-EM structure of PIEZO1 (PDB: 5Z10), PIEZO1-curved (PDB: 7WLT) and PIEZO1-flattened (PDB:7WLU) were aligned to the cryo-EM map by Chimera, and used as the initial model, respectively. Manual fitting was performed using COOT^61^. Then the model was refined in real space using PHENIX with secondary structure and non-crystallographic symmetry (NCS) restraints. Lipid-like densities that appeared in the density maps were fitted by trial with candidate lipids. The final atomic model was validated using MolProbity^62^.

### Measurement of the curvature radius of the R2482H proteoliposome

To measure the curvature variance induced by the reconstituted PIEZO1-R2482H in proteoliposome vesicles, the distance between the central point of three inner helices and the center site of the bilayer was measured by IMOD^63^ as r_project_ (radius of the open mouth plane of the bowl). The projected area of the membrane is derived from the formula for the area of a circle: A_project_ = πr_project_². As described above (Fig. 2), the nano-bowl shape of the R2482H-liposome-curved has a depth of 12.5 nm, giving rise to an in-plane projection area of 490.9 nm^2^. The co-planar PIEZO1 membrane of the R2482H-liposome-flattened has a radius of 15 nm and an area of 706.5 nm^2^.

### HOLE and CHAP analyses

The central solvent-accessible pathway is generated by the program HOLE^64^. CHAP^65^ analysis was performed according to online instructions (https://www.channotation.org/). The PDB files of four structures were used as input for CHAP analysis. The output files of this analysis include the time-averaged hydrophobicity due to pore-facing residues and pore radii.

### Whole-cell electrophysiology

The whole-cell configuration was used for measuring the poking-evoked currents as previously described^33^. The electrophysiological recordings were carried out with HEKA EPC10 at room temperature. The recording electrodes (Sutter Instrument) had a resistance of 2-6 MΩ when filled with internal buffer solution (in mM: 133 CsCl, 1 MgCl2, 1 CaCl2, 5 EGTA, 10 HEPES, pH 7.3 with CsOH, 4 MgATP and 0.4 Na2GTP). The extracellular bath solution was (in mM) 133 NaCl, 1 MgCl2, 2.5 CaCl2, 3 KCl, 10 HEPES (pH 7.3 with NaOH) and 10 glucose. Data were sampled at 20 kHz and filtered at 2.9 kHz by Patchmaster software. In offline analysis, currents were further filtered at 1 kHz and leak currents before mechanical stimulation were subtracted. The inactivation time of maximal current per cell was fitted by a mono-exponential equation.

A fire-polished glass pipette (tip diameter 3-4 μm) was used to indent cells by an amplifier-controlled piezo-electric driver (E625 LVPZT Controller/Amplifier; Physik Instrumente). The probe was positioned close to the cell at an angle of 80° and advanced at 1 μm/ms during the ramp segment of the downward and upward motion. For measure inactivation, a stimulus sufficient to elicit a robust current was applied and maintained for 150 ms. The time course of inactivation was fitted with a single exponential function. The deactivation was measured as previously described. In brief, the stimulus was promptly withdrawn following its application. For inward currents, deactivation kinetics were fitted with a single-exponential function, whereas outward currents required a double-exponential function. Fitting was performed starting from an inflection point in the current decay, which likely represents the onset of channel deactivation during retraction of the poking probe. An inter-stimulus interval of 8 s was used for both protocols.

### Cell-attached electrophysiology

The cell-attached configuration was used for recording single-channel events as previously described^33^. All experiments were carried out with HEKA EPC10 at room temperature. The recording electrodes (Sutter Instrument) had a resistance of 2-6 MΩ when filled with internal buffer solution (in mM: 130 NaCl, 5 KCl, 1 CaCl2, 1 MgCl2, 10 TEA-Cl and 10 HEPES, pH 7.3 with NaOH). The extracellular bath solution consisted of (in mM) 140 KCl, 1 MgCl2, 10 glucose and 10 HEPES (pH 7.3 with KOH). Membrane patches with a seal resistance of at least 2 GΩ were stimulated with negative pressure pulses using a high-speed pressure clamp system (HSPC-1; ALA-scientific). Data were sampled at 20 kHz by Patchmaster software. In offline analysis, currents were further filtered at 1 kHz, and leak currents before mechanical stimulation were subtracted. Cells were held at -120 mV for recording single-channel currents. For each potential, 70 to 120 repeated recordings were performed. For calculating single-channel currents, the Gaussian equation was used to fit the current histogram in the Clampfit 10 software.

### Measurement of permeability ratios

To measure the chloride-to-sodium permeability ratio (P_Cl_/PNa), the internal solution consisted of (in mM) 150 mM NaCl and 10 mM HEPES (pH 7.3 with NaOH). The external solution contained (in mM) 30 mM NaCl, 10 mM HEPES, and 225 mM sucrose (pH 7.3 with NaOH). The reversal potential (Erev) for each cell was measured using linear regression fit to the I-V relationship curve under the corresponding solution conditions. The permeability ratios were calculated using the following Goldman– Hodgkin–Katz (GHK) equation:

For the permeability ratio of Cl^−^ to Na^+^ (P_Cl_/P_Na_):

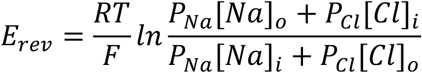

For the permeability ratio of Ca^2+^ to Cs^+^ (P_Ca_/P_Cs_):

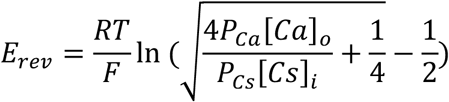

Where 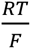 equals 25.69 at 25°C.

## Extended Data Figures and Legends

**Extended Data Fig. 1.**
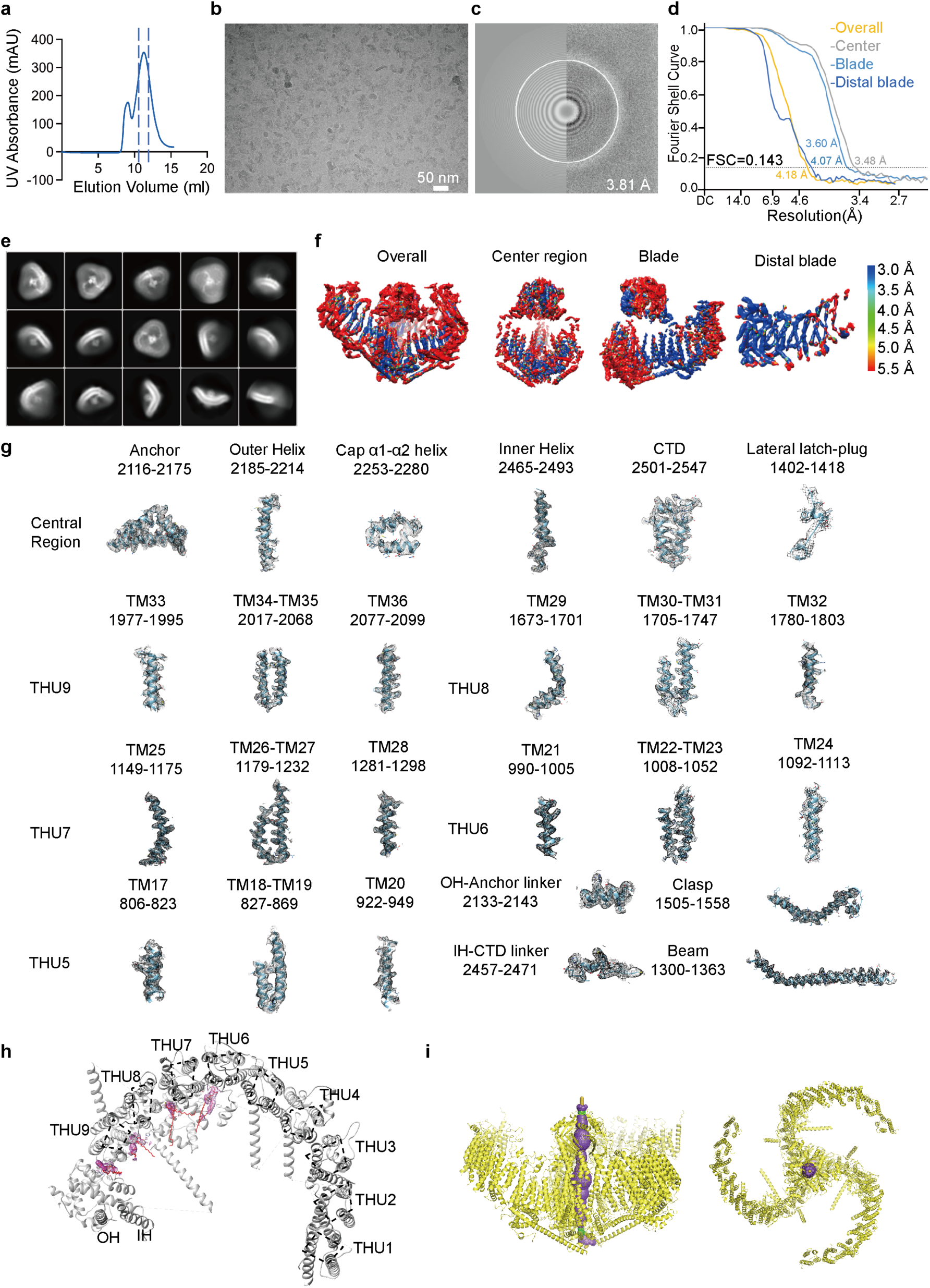
Cryo-EM determination of the R2482H-detergent-curved structure. **a**, A representative trace of gel filtration of purified PIEZO1-R2482H proteins. UV, ultraviolet. Dash lines indicate the peak fractions collected for cryo-sample preparation. **b**, A representative cryo-electron micrograph of PIEZO1-R2482H solubilized in the detergent GDN. **c**, Power spectrum of the micrograph in b, with the 3.81 Å frequency indicated. **d**, Gold-standard Fourier shell correlation (FSC) curves of center, peripheral blade and overall density maps. Reported resolutions were based on the FSC = 0.143 criterium. **e**, Representative 2D class averages. **f**, Local-resolution subtracted maps. **g**, Local EM density of the indicated domains. The helices are shown as cartoons with side chains as sticks, fitting with the cryo-EM density shown as gray mesh. **h**, A top view of a single blade of the PIEZO1-R2482H structure with resolved 38-TMs, the lipid densities flanking the blade were highlighted in magenta. **i**, The side (left) and top (right) views of the PIEZO1-R2482H structure cartoon model. The pore region was highlighted as the purple mesh.

**Extended Data Fig. 2.**
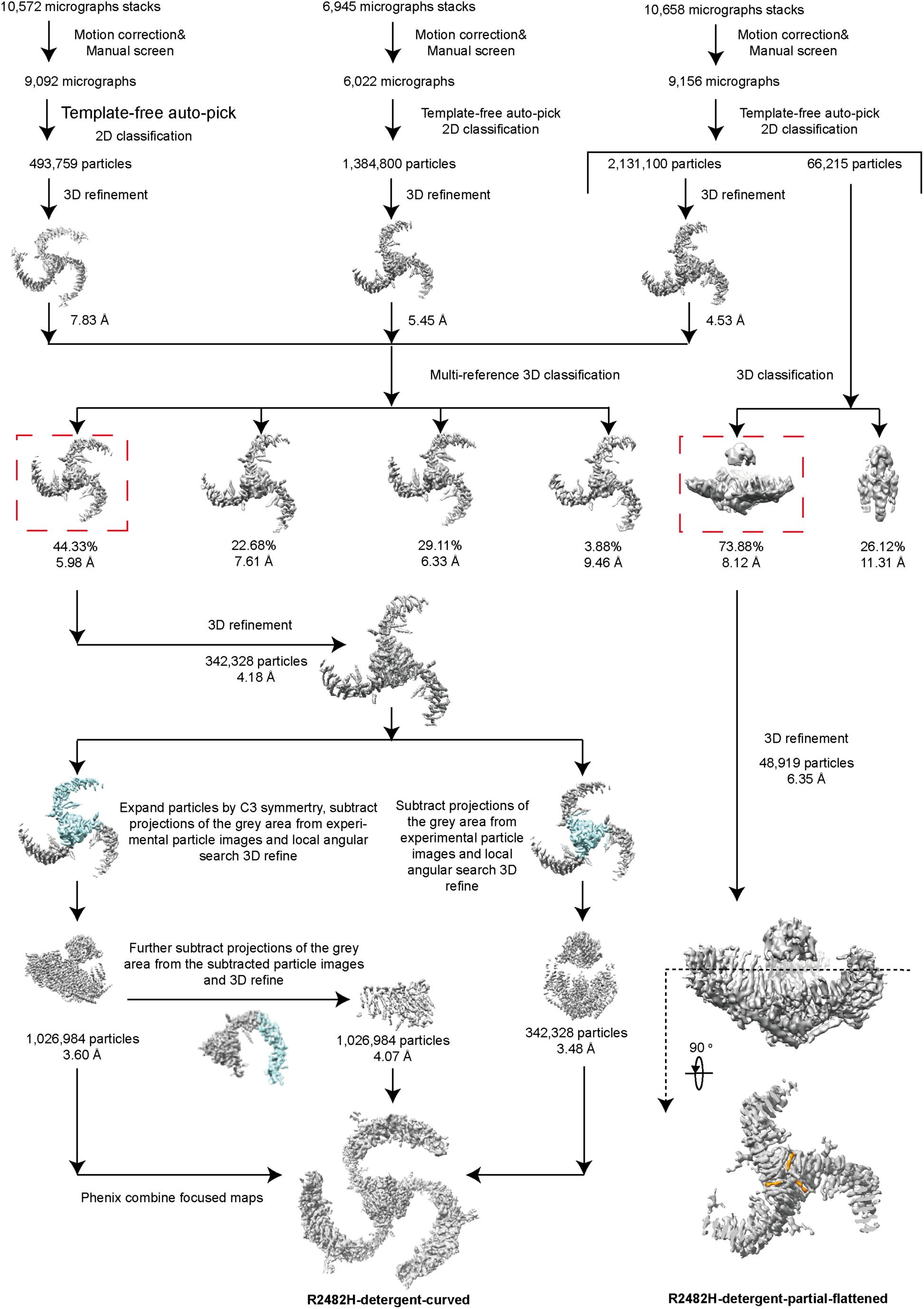
Flowchart of electron-microscopy data processing of the R2482H-detergent-curved structure. Details of data processing are described in the ‘Imaging processing’ section of the Methods.

**Extended Data Fig. 3.**
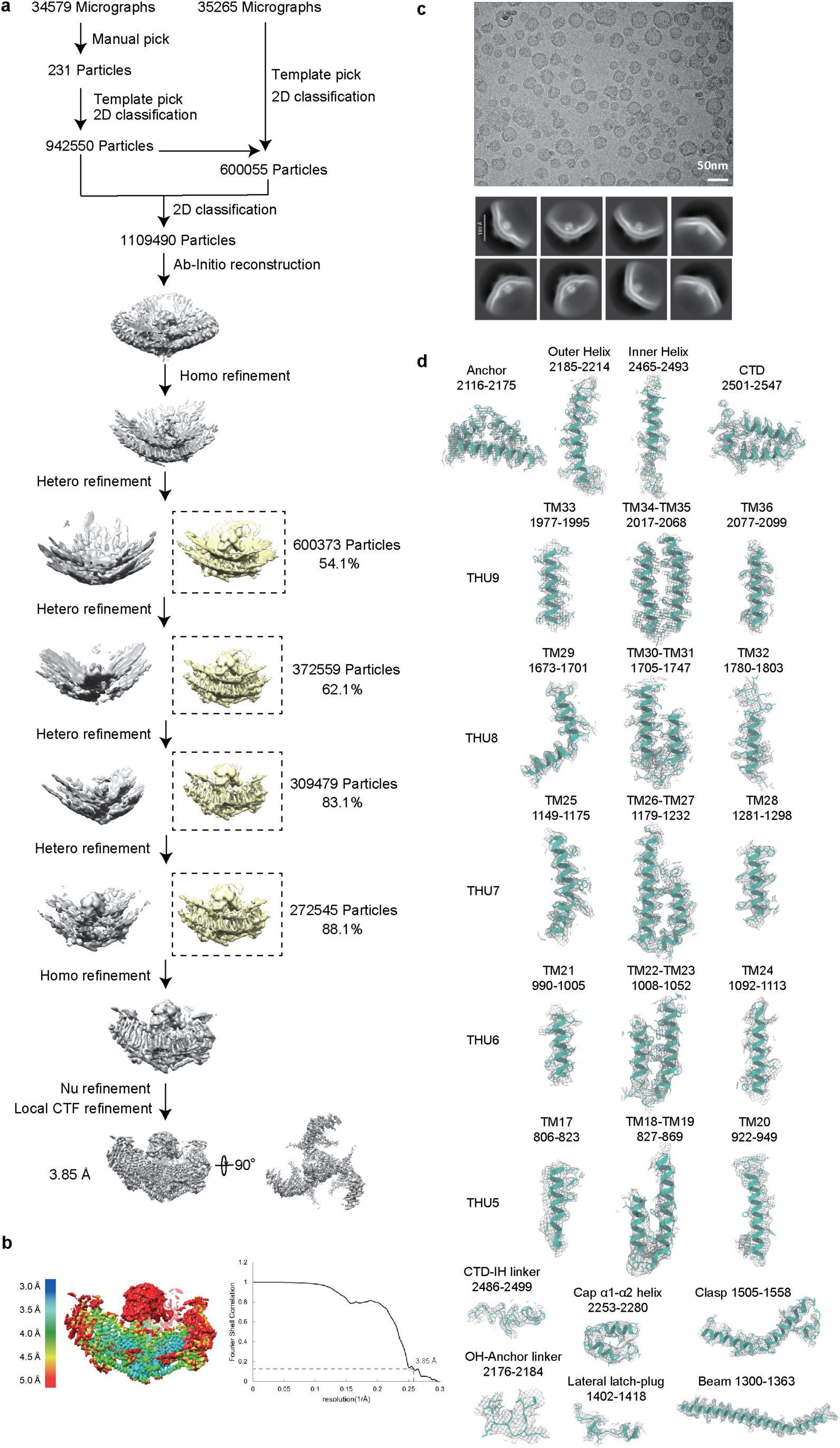
Cryo-EM determination of the R2482H-liposome-curved structure in proteoliposomes. **a,** Flowchart of electron-microscopy data processing of the R2482H-liposome-curved structure. **b,** Local-resolution map and gold-standard Fourier shell correlation (FSC) curve of the R2482H-liposome-curved structure. Reported resolutions were based on the FSC = 0.143 criterium. **c,** A representitive cryo-electron micrograph of R2482H proteoliposome, and representative 2D classes of R2482H-liposome-curved. **d,** Local EM density of the R2482H-liposome-curved structure. The helices are shown as cartoons, fitting with the cryo-EM density shown as the gray mesh.

**Extended Data Fig. 4.**
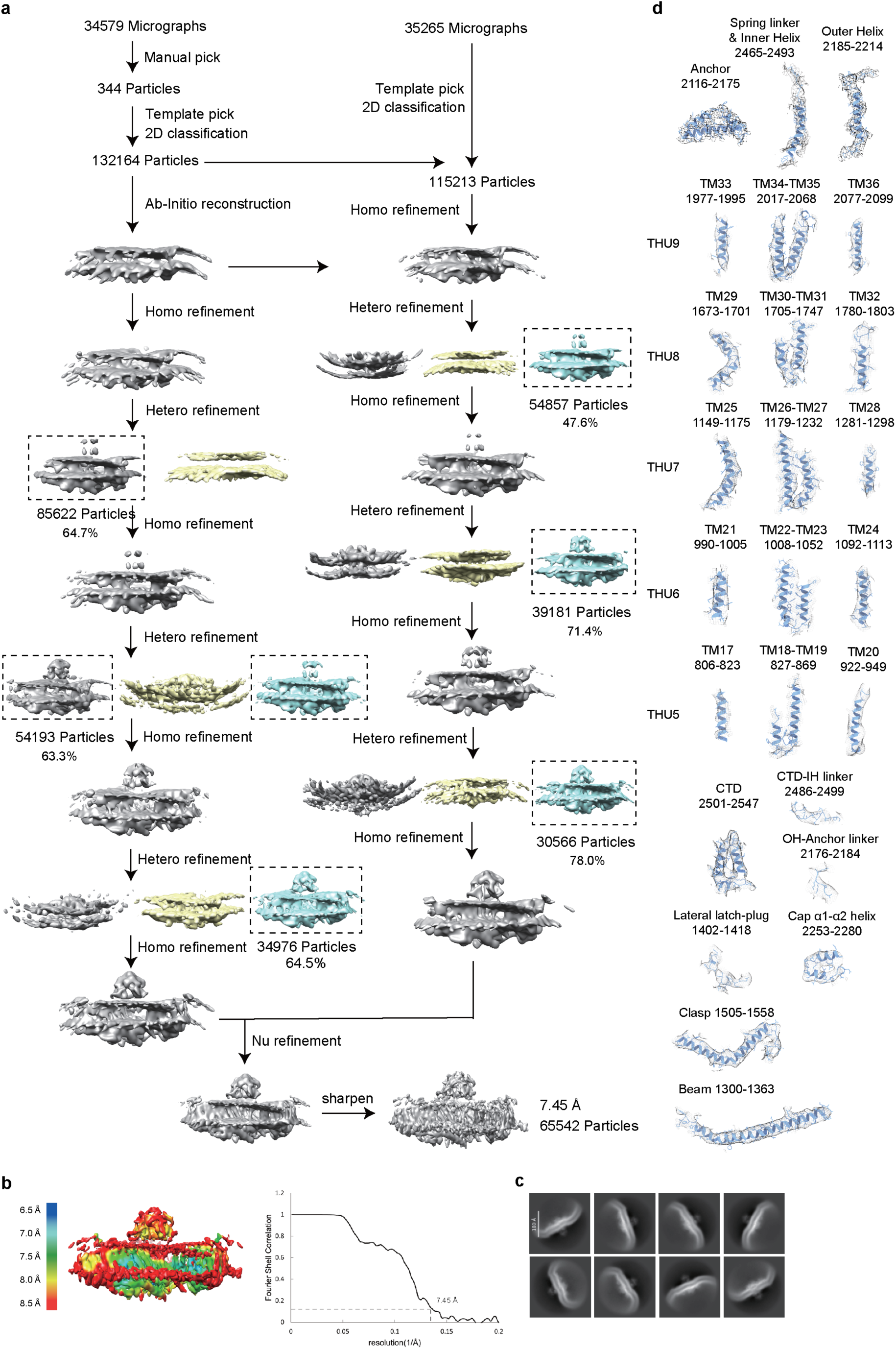
Cryo-EM determination of the R2482H-liposome-flattened structure in proteoliposomes. **a,** Flowchart of electron-microscopy data processing of the R2482H-liposome-flattened structure. **b,** Local-resolution map and gold-standard Fourier shell correlation (FSC) curve of the R2482H-liposome-flattened structure. Reported resolutions were based on the FSC = 0.143 criterium. **c,** Representative 2D classes of R2482H-liposome-flattened. **d,** Local EM density of the R2482H-liposome-flattened structure. The helices are shown as cartoons, fitting with the cryo-EM density shown as the gray mesh.

**Extended Data Fig. 5.**
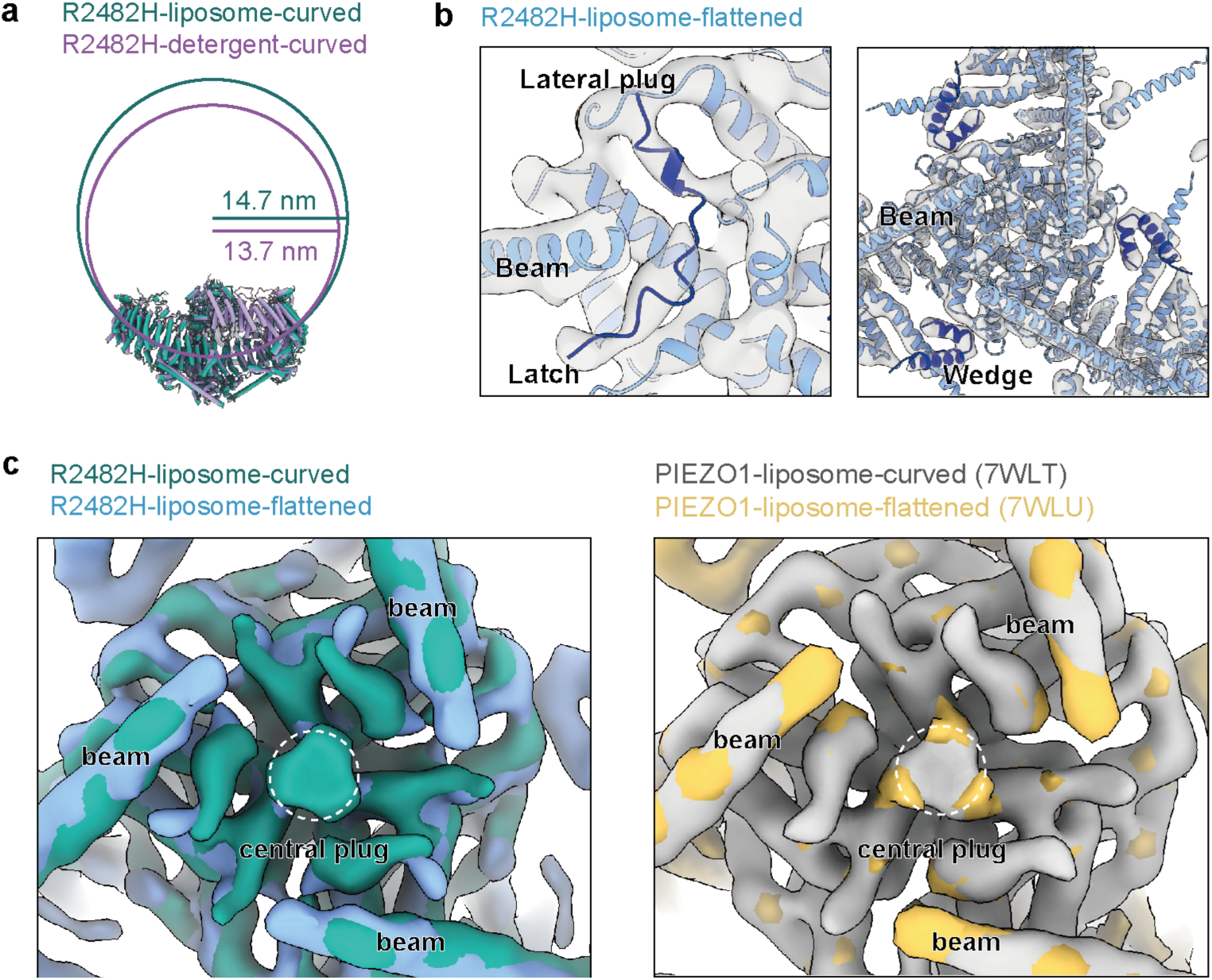
Structural analyses of R2482H-liposome-curved and R2482H-liposome-flattened. **a,** Curvature difference between the R2482H-detergent-curved and R2482H-liposome-curved structures. The measured geometric parameters are listed in Extended Data Table 2. **b,** A transparent view of the cryo-EM density map showing fragmented density of the lateral-plug and latch domains and the folded wedge domain. **c,** Bottom view of the overlaid curved and flattened maps of either R2482H-liposome and PIEZO1-liposome showing the disassembly of central plug density in the flattened maps.

**Extended Data Fig. 6.**
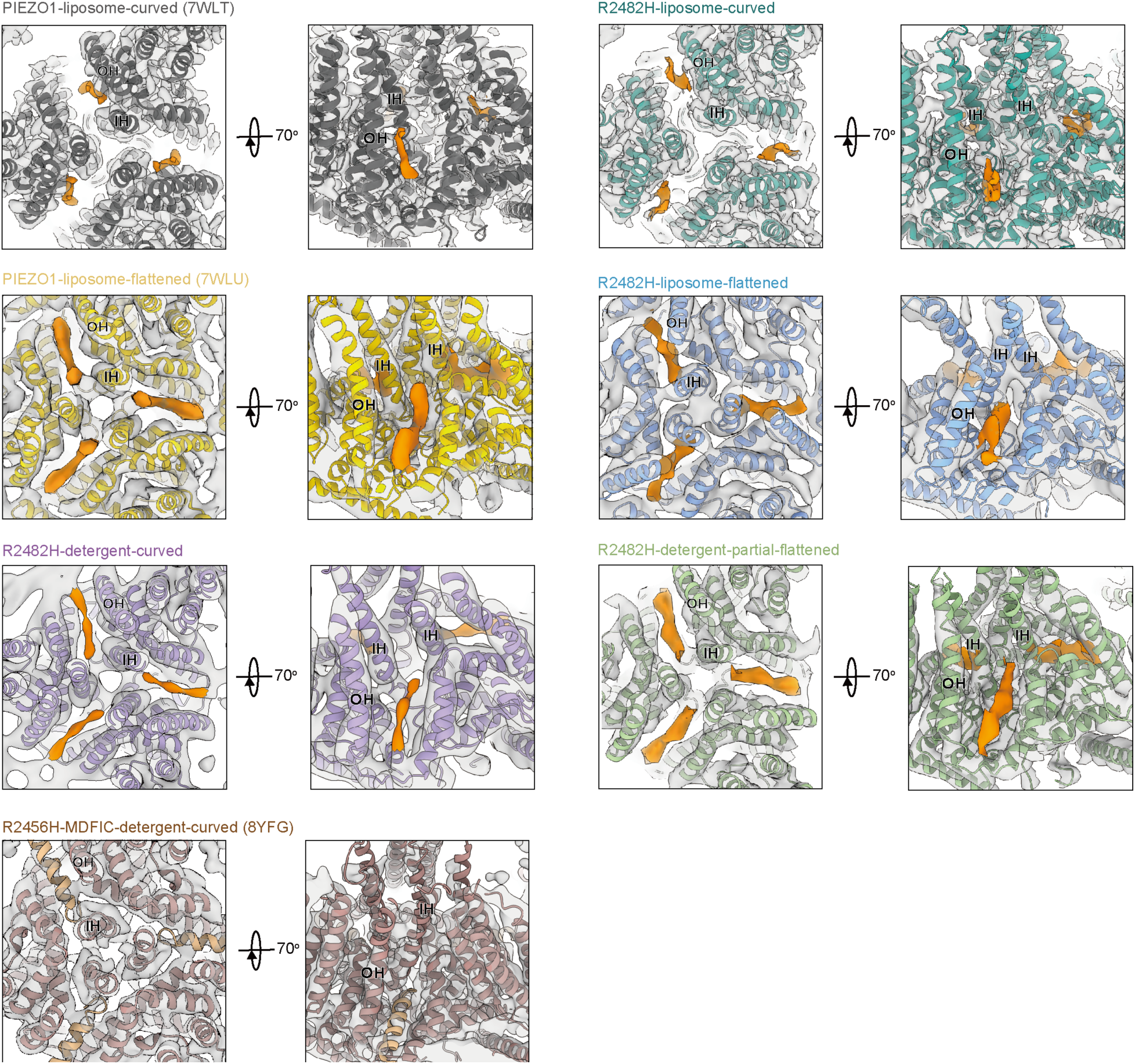
Lipid-like density in the lateral portals of the PIEZO1 and R2482H structures. Top-down and side views of the central pore module of the indicated structures showing the lipid-like density (in orange) right above the lateral portal. The densitiy map of R2482H-detergent-curved is low-pass filtered to 8 Å.

**Extended Data Fig. 7.**
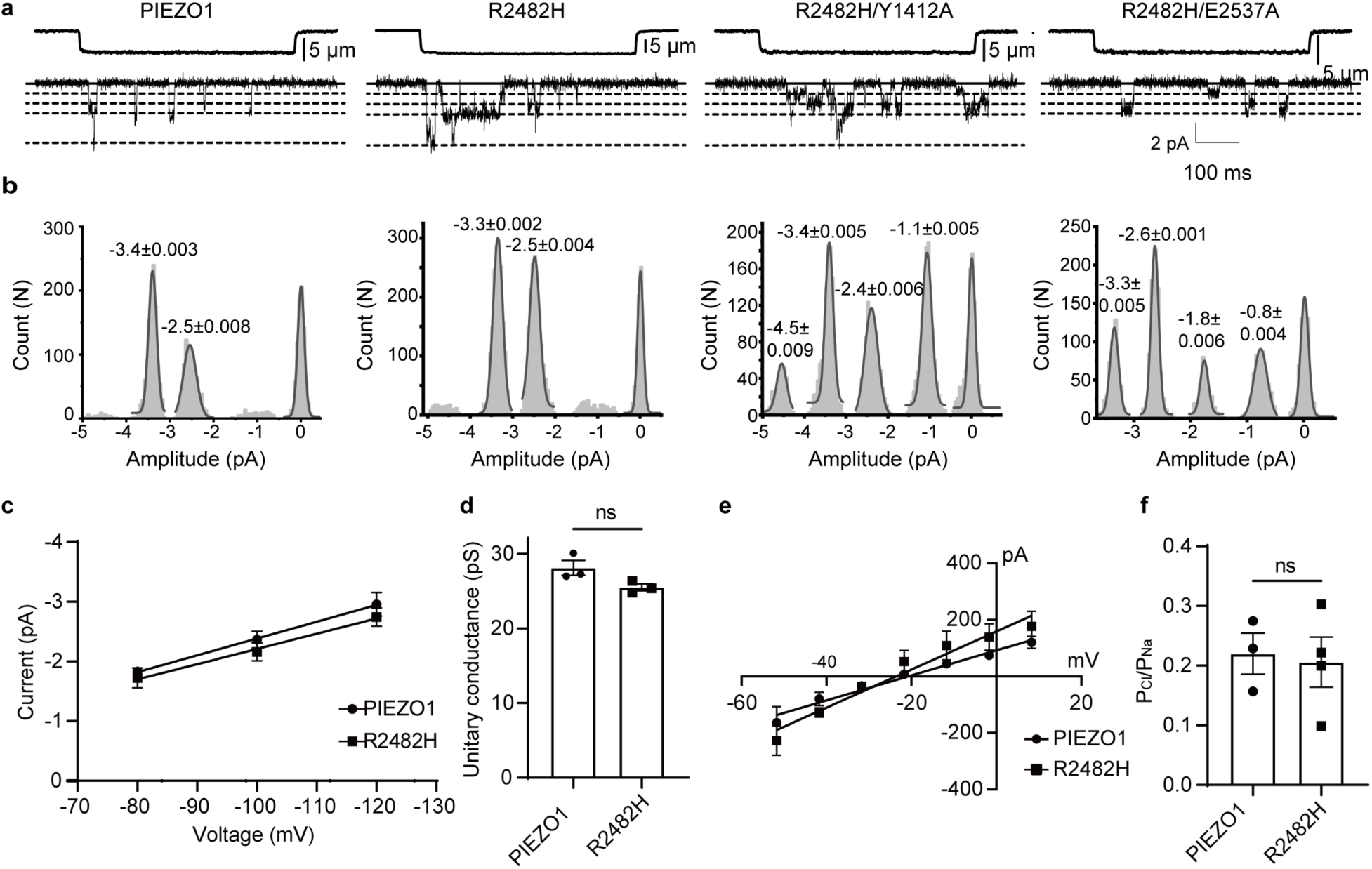
R2482H does not affect the ion selectivity and permeation properties of PIEZO1. **a**, Representative traces of single-channel currents from cells transfected with the indicated constructs at -120 mV. The dashed lines indicate 1/3, 2/3 and 3/3 of the full single-channel conductance. **b**, Histogram analysis of the single-channel currents of the indicated constructs from (**a**). The peak currents are labeled. **c**, Average I-V relationships of stretch-induced single-channel currents in HEK293T cells transfected with the indicated constructs (mean ± SEM, n=3). **d**, The unitary conductance of the indicated constructs (mean ± SEM, n=3; Unpaired Student’s t-test; ns represents non-significance). **e**, Average I-V relationships of the mechanically evoked whole-cell currents mediated by PIEZO1 or R2482H indicated constructs for determining the permeability ratios of P_Cl_/P_Na_ (mean ± SEM, n=3-4). **f**, The P_Cl_/P_Na_ permeability ratio derived from poking-activated currents mediated by PIEZO1 or R2482H (mean ± SEM, n=3-4; Unpaired Student’s t-test; ns represents non-significance).

**Extended Data Table 1.** Cryo-EM data collection, refinement and validation <u>statistics</u>.

|  | R2482H-detergent-curved<br>(EMD-xxxxx)<br>(PDB xxxx) | R2482H-liposome-curved<br>(EMD-xxxxx)<br>(PDB xxxx) | R2482H-liposome-flattened<br>(EMD-xxxxx)<br>(PDB xxxx) |
| --- | --- | --- | --- |
| **Data collection and processing** |  |  |  |
| Magnification | 64,000 | 64,000 | 64,000 |
| Voltage (kV) | 300 | 300 | 300 |
| Electron exposure(e<sup>-</sup>/Å<sup>2</sup>) | 50 | 50 | 50 |
| Defocus range (μm) | -1.5~-2.0 | -1.3~-1.8 | -1.3~-1.8 |
| Pixel size (Å) | 1.098 | 1.098 | 1.098 |
| Symmetry imposed | C3 | C3 | C3 |
| Initial particle images (no.) | 1,026,984 | 1,109,490 | 247,377 |
| Final particle images (no.) | 342,328 | 272,545 | 65,542 |
| Map resolution (Å) | 4.18 | 3.85 | 7.45 |
| FSC threshold | 0.143 | 0.143 | 0.143 |
| Map resolution range (Å) | 5.9-3.0 | 5.0-3.0 | 8.5-6.5 |
| **Refinement** |  |  |  |
| Initial model used (PDB code) | 5Z10 | 7WLT | 7WLU |
| Model resolution (Å) | 4.3 | 4 | 7.6 |
| FSC threshold | 0.5 | 0.5 | 0.5 |
| Map sharpening *B* factor (Å<sup>2</sup>) | -105 | -147 | -268 |
| Model composition |  |  |  |
| Non-hydrogen atoms | 45433 | 37605 | 34374 |
| Protein residues | 5544 | 4512 | 4458 |
| Ligands | 27 | 27 | 0 |
| *B* factors (Å<sup>2</sup>) |  |  |  |
| Protein | 142.82 | 108.87 | 327.19 |
| Ligand | 106.88 | 103.77 |  |
| R.m.s. deviations |  |  |  |
| Bond lengths (Å) | 0.006 | 0.006 | 0.002 |
| Bond angles (°) | 0.988 | 0.809 | 0.698 |
| Validation |  |  |  |
| MolProbity score | 2.29 | 2.37 | 2.08 |
| Clashscore | 27.62 | 11.80 | 12.02 |
| Poor rotamers (%) | 0.00 | 0.00 | 0.00 |
| Ramachandran plot |  |  |  |
| Favored (%) | 94.83 | 95.20 | 92.04 |
| Allowed (%) | 5.06 | 4.78 | 7.82 |
| Disallowed (%) | 0.11 | 0.02 | 0.14 |

**Extended Data Table 2.** Geometric parameters of PIEZO structures.

| PIEZO structures | The outmost TM | r (Å) | h (Å) | R (Å) |
| --- | --- | --- | --- | --- |
| PIEZO2-detergent-curved (6KG7) | TM3 of THU1 | 114 | 91 | 117 |
| PIEZO2-detergent-curved (6KG7) | TM15 of THU4 | 105 | 66 | 116 |
| PIEZO1-detergent-curved (5Z10) | TM15 of THU4 | 98 | 52 | 118 |
| PIEZO1-detergent-curved (6B3R) | TM15 of THU4 | 101 | 52 | 123 |
| PIEZO1-liposome-curved (7WLT) | TM19 of THU5 | 90 | 52 | 103 |
| PIEZO1-liposome-flattened (7WLU) | TM15 of THU4 | 120 | 7 | 1091 |
| S2472E-detergent-curved | TM15 of THU4 | 108 | 53 | 137 |
| S2472E-detergent-intermediate-open | TM15 of THU4 | 123 | 25 | 318 |
| R2482H-detergent-curved | TM3 of THU1 | 112 | 58 | 137 |
| R2482H-detergent-curved | TM15 of THU4 | 106 | 50 | 137 |
| R2482H-detergent-partial-flattened | TM15 of THU4 | 113 | 44 | 168 |
| R2482H-liposome-curved | TM15 of THU4 | 108 | 47 | 147 |
| R2482H-liposome-flattened | TM15 of THU4 | 123 | 3 | 2784 |

---

**Video S1.** Structural rearrangement and gating cycle of PIEZO1.

**Video S2.** Structural rearrangement and gating cycle of the central pore of PIEZO1.

**Video S3.** Structural rearrangement and gating cycle of R2482H.

**Video S4.** Structural rearrangement and gating cycle of the central pore of R2482H.

## Notes

### Competing Interest Statement

The authors have declared no competing interest.

